# Cdc6 is sequentially regulated by PP2A-Cdc55, Cdc14 and Sic1 for origin licensing in *S. cerevisiae*

**DOI:** 10.1101/2021.09.14.460335

**Authors:** Jasmin Philip, Mihkel Örd, Andriele Silva, Shaneen Singh, John F. X. Diffley, Dirk Remus, Mart Loog, Amy Ikui

## Abstract

Cdc6, a subunit of the pre-replicative complex, contains multiple regulatory Cdk1 consensus sites, SP or TP motifs. In *S. cerevisiae*, Cdk1 phosphorylates Cdc6-T7 to recruit Cks1, the Cdk1 phospho-adaptor in S-phase, for subsequent multisite phosphorylation and protein degradation. Cdc6 accumulates in mitosis and is tightly bound by Clb2 through N-terminal phosphorylation in order to prevent premature origin licensing and degradation. It has been extensively studied how Cdc6 phosphorylation is regulated by the Cyclin-Cdk1 complex. However, a detailed mechanism on how Cdc6 phosphorylation is reversed by phosphatases has not been elucidated. Here, we show that PP2A^Cdc55^ dephosphorylates Cdc6 N-terminal sites to release Clb2. Cdc14 dephosphorylates the C-terminal phospho-degron, leading to Cdc6 stabilization in mitosis. In addition, the Cdk1 inhibitor, Sic1, releases Clb2·Cdk1·Cks1 from Cdc6 to load Mcm2-7 on the chromatin upon mitotic exit. Thus, pre-RC assembly and origin licensing is promoted by the attenuation of distinct CDK-dependent Cdc6 inhibitory mechanisms.

## Introduction

Pre-replicative complexes (pre-RCs) are assembled on DNA to license replication origins in M-G1 phase. The pre-RC components are recruited in a sequential fashion that starts with the origin of replication complex (Orc1-6) followed by Cdc6 then Cdt1, which eventually load the Mcm2-7 helicase on DNA (Bell & Stillman, 1992; Labib *et al*., 2001; Newlon, 1997; Santocanale & Diffley, 1996; Tanaka & Diffley, 2002). At the onset of S-phase, cyclin-dependent-kinase (Cdk1) phosphorylates pre-RC components such as Cdc6, which prevents re-initiation of DNA replication through multiple mechanisms (Nguyen *et al*., 2001; Wilmes *et al*., 2004). Cdc6 phospho-degrons are targeted by CDK, which directs Cdc6 protein degradation via SCF-mediated ubiquitination in *S. cerevisiae* (Drury *et al*., 1997; Drury *et al*., 2000; Perkins *et al*., 2001). Cdc6 is expressed and stabilized during late mitosis; however, it is not clear how Cdc6 stability is maintained in the presence of high Cdk1 activity.

There are eight Cdk1-dependent phosphorylation sites that contain TP or SP motifs, six of which have been well characterized (Figure 1A). Cdk1-mediated phosphorylation is facilitated by specific cyclin docking motifs in substrates: RxL cyclin motif for Clb5-Cdk1 interaction and LxF motif for Clb2-Cdk1 (Cross & Jacobson, 2000; Kõivomägi *et al*., 2011; Loog & Morgan, 2005; Örd *et al*., 2019; Wilmes *et al*., 2004). S-phase cyclin Clb5 binds to the Cdc6 RxL motif at amino acid residues 29-31 to mediate phosphorylation of T7 in the Cdc6 N-terminus, which serves as a Cks1 docking site. The docking event triggers Cdc6 multi-phosphorylation cascade in N- to C-direction (Örd *et al*., 2019). SCF^Cdc4^ recognizes Cdc6 phospho-degrons at T39-S43 and T368-S372 for ubiquitin-mediated degradation under control of the SCF^Cdc4^ ubiquitin ligase (Figure 1A) (Al-Zain *et al*., 2015; Drury *et al*., 1997; Drury *et al*., 2000; Perkins *et al*., 2001).

**Figure 1.**
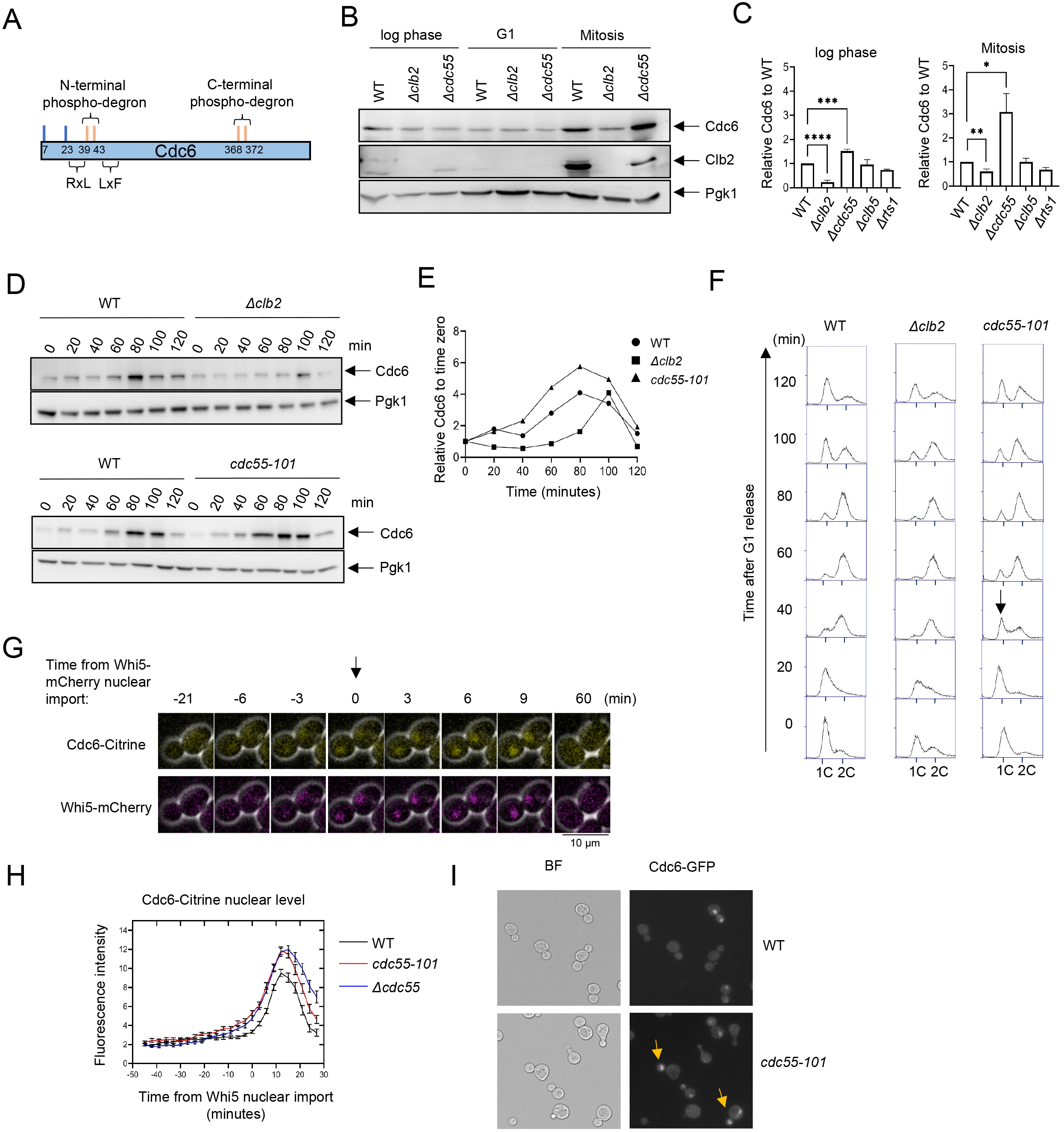
Cdc6 is more stabilized in *Δcdc55* cells than in *Δclb2*. A) Cdc6 contains six functional Cdk1 phosphorylation sites including N-terminal and C-terminal phospho-degrons at S39-T43 and S368-T372, respectively (orange bars). RxL and LxF motifs mediate cyclin interaction. B) *CDC6-9MYC* (WT), *CDC6-9MYC Δclb2* (*Δclb2*) or *CDC6-9MYC Δcdc55* (*Δcdc55*) cells were incubated to log-phase, or arrested in G1 by alpha-factor or mitosis by nocodazole. Protein was extracted and subjected to western blot analysis to visualize Cdc6-9MYC and Clb2. Pgk1 was used as a loading control. C) Cdc6 protein levels from A were quantified and normalized to WT. Three independent biological replicates were performed. The average of relative Cdc6 band intensity compared to WT is shown. Error bar is SEM (n=4; log phase *Δclb2 \*\*\*\**p<0.0001, *Δcdc55 \*\*\**p=0.0006, Mitosis *Δclb2 \*\**p=0.0075, *Δcdc55 \**p=0.0351 was calculated by unpaired Student t-test) D) *CDC6-PrA* (WT), *CDC6-PrA Dclb2* (*Δclb2*), *CDC6-PrA cdc55-101 (cdc55-101)* were synchronized in G1 phase by alpha-factor and released. Samples were collected at indicated times. Cdc6-PrA was visualized by western blotting analysis. Pgk1 was used as a loading control. E) The relative Cdc6 band intensity to time zero from C is shown. F) Samples from C were fixed and stained with propidium iodide (PI) to show cell cycle profile by flow cytometry. X-axis is PI and Y-axis is cell number. Arrow indicates S-phase entry delay. G) *CDC6-Citrine WHI5-mCherry* cells were imaged by time-lapse microscopy. Time zero was set when Whi5-mCherry is imported to the nucleus. Arrow marks Whi5 nuclear entry in late mitosis. H) *CDC6-Citrine* (WT), *CDC6-Citrine cdc55-101* (*cdc55-101)* or *CDC6-Citrine Dcdc55* (*Δcdc55*) with Whi5-mCherry were imaged by time-lapse microscopy. Shown is the average Cdc6-Citrine fluorescence intensities of 100 cells per time point. Error bars are SEM (n=100). I) *CDC6-GFP* (WT) or *CDC6-GFP cdc55-101* (*cdc55-101)* cells were grown to log-phase and imaged using a fluorescence microscope.

Following Cdc6 degradation at G1/S, it is expressed again in mitosis and is tightly bound to mitotic cyclin Clb2 to prevent origin licensing in mitosis (Mimura *et al*., 2004). Clb2 interacts with Cdc6 through the Clb2 hydrophobic patch (*hp*), a substrate-binding pocket in cyclins (Örd *et al*., 2019). The specific binding of Clb2 to Cdc6 is mediated through the ^47^LQF^49^ motif at the Cdc6 N-terminus and is enhanced by a ^126^FQSLP^130^ motif located in the mid-region of Cdc6 (Örd *et al*., 2019). Thus, the tight Clb2-Cdc6 interaction potentially shields Cdc6 phospho-degrons, leading to Cdc6 stabilization. These mechanisms could accumulate the Cdc6 protein in mitosis without origin licensing.

Mitotic exit requires a complete suppression of Cdk1 activity to reset the cell cycle for G1 entry. It has been suggested that Cdc6 acts as a CDK inhibitor together with Sic1 and Cdh1, a co-activator of the APC (Calzada *et al*., 2001). Sic1 and Cdc6 share similarities in Cdk1 inhibitory function, as both have been shown to associate cyclin-Cdk1 complexes with low nanomolar affinity which is partly facilitated by Cks1 docking (Örd *et al*., 2019; Venta *et al*., 2020). Deletion of the Cdc6 N-terminal Cdk1 site (*cdc6Δ47)* shows an elevated Clb2-Cdk1 activity in mitosis, indicating that Cdc6 N-terminus plays a role in Clb2-Cdk1 inhibition (Calzada *et al*., 2001). A combination of *Δsic1* and *cdc6Δ47* leads to defects in mitotic exit, probably due to high Cdk1 activity (Calzada *et al*., 2001). Furthermore, *Δsic1 Δcdh1 cdc6Δ2-49* triple mutants caused cytokinesis defects (Archambault *et al*., 2003). Altogether, Cdc6, Sic1, and APC^Cdh1^ cooperate to inhibit Clb2-Cdk1 activity for mitotic exit. This was further supported by an *in vitro* assay that shows the inhibition of Clb2/Cdc28 kinase activity by Cdc6 (Örd *et al*., 2019).

Beside these Cdk1 inhibition mechanisms, Cdc6 phosphorylation needs to be removed by phosphatases during late mitosis. It is unknown which phosphatase is responsible for Cdc6 dephosphorylation. It has been reported that Cdc6 physically interacts with Cdc55, a regulatory subunit for protein phosphatase 2A (PP2A) (Boronat & Campbell, 2007). PP2A belongs to a family of serine/threonine phosphatases with a well-conserved role in mitosis from yeast to humans (Janssens & Goris, 2001; Mochida *et al*., 2009). In *S. cerevisiae*, the PP2A heterotrimeric complex is composed of a scaffold A subunit (Tpd3), a regulatory B subunit (Cdc55, Rts1 or Rts3), and two largely interchangeable catalytic C subunits (Pph21 and Pph22) (Healy *et al*., 1991; Shu *et al*., 1997; van Zyl *et al*., 1992). Among these subunits, the regulatory B subunit primarily dictates cellular localization and PP2A substrate specificity (Rossio & Yoshida, 2011).

PP2A^Cdc55^ and PP2A^Rts1^ have been implicated in mitotic progression. Cytoplasmic PP2A^Cdc55^ dephosphorylates and inhibits Swe1, a Cdk1 inhibitor, to drive mitotic entry (F. C. Lin & Arndt, 1995; Yang *et al*., 2000). Nuclear PP2A^Cdc55^ plays a role in the spindle assembly checkpoint and inhibits chromosome segregation by dephosphorylating Cdc20 (Lianga *et al*., 2013; Minshull *et al*., 1996; Rossio *et al*., 2013; Wang & Burke, 1997). Furthermore, various stresses inhibit the cell cycle via PP2A^Cdc55^ (Khondker *et al*., 2020; Tang & Wang, 2006). PP2A^Cdc55^ also dephosphorylates Net1, an inhibitor of Cdc14 (Queralt *et al*., 2006). PP2A^Rts1^ promotes cytokinesis by modulating actin ring phosphorylation (Dobbelaere *et al*., 2003). It is unclear if PP2A directly targets and dephosphorylates Cdc6.

Cdc14 phosphatase promotes mitotic exit through Net1 dephosphorylation upon Cdc14 release from the nucleolus known as FEAR and MEN networks (Azzam *et al*., 2004; Shou *et al*., 1999; Tomson *et al*., 2009; Visintin *et al*., 1999). Cdc14 dephosphorylates cell cycle regulators such as Swi5, Sic1 and Cdh1, all which are necessary for mitotic exit and G1 entry (Jaspersen *et al*., 1999; Visintin *et al*., 1998). It has been reported that Cdc6 is destabilized in *cdc14-3* mutant, which prompted us to study if Cdc14 directly dephosphorylates Cdc6 (Zhai *et al*., 2010). In this study, we show that Cdc6 is directly dephosphorylated by PP2A^Cdc55^ and Cdc14 at distinct Cdk1 sites in the Cdc6, resulting in Clb2 release from the inhibitory complex. Sic1 also releases Clb2 from Cdc6, allowing Mcm2-7 loading on chromatin for pre-RC assembly. These results suggest a model in which distinct phosphatases relieve CDK-dependent Cdc6 inhibition. Finally, a structural insight into Cdc6-Clb2 binding using bioinformatics suggests a role of the Cdc6 intrinsically disordered region for multiple protein-protein interactions.

## Results

### Clb2 and PP2A^Cdc55^ have contrasting roles in Cdc6 stability

To test the idea that PP2A^Cdc55^ targets Cdc6, we examined Cdc6 protein levels in *Δclb2* and *Δcdc55* cells by Western blot analysis. The Cdc6 protein level was lower in *Δclb2* compared to wild type in log phase cells, which supports previous results of Cdc6 destabilization in the Cdc6*-*lxf mutant that does not bind to Clb2 (Figure 1B left and 1C) (Örd *et al*., 2019). There was low Cdc6 protein expression in G1 arrested cells which was not impacted by deletion of *CLB2* or *CDC55* (Figure 1B, middle). During mitosis, we observed less Cdc6 in *Δclb2* cells and more in *Δcdc55* cells compared to wild type; this difference in the Cdc6 protein level was more prominent in mitosis than in log-phase cells (Figure 1B right and 1C). These results indicate that PP2A^Cdc55^ and Clb2 regulate Cdc6 protein levels in a contradictory manner. An elevated Clb2 protein level and Cdc6 stabilization in wild type cells confirmed that cells were in mitosis (Figure 1B right). The S-phase cyclin Clb5, as well as another regulatory subunit of PP2A, Rts1, did not alter Cdc6 protein level (Figure 1C). We conclude that Cdc6 is regulated by mitotic cyclin Clb2 and PP2A^Cdc55^ in a specific manner.

Next, we analyzed Cdc6 protein levels in *Δclb2* and *cdc55-101* cells during the cell cycle. The *cdc55-101* mutant excludes Cdc55 from the nucleus, allowing us to study Cdc55 nuclear function. Unlike *Δcdc55*, *cdc55-101* mutant cells do not exhibit morphological defects, which allows us to monitor G1 arrest induced by α-factor and cell cycle progression (Sasaki *et al*., 2000). In wild type cells, Cdc6 protein levels were suppressed throughout S-phase, increased in mitosis at 80 minutes and then diminished (Figure 1D and 1E) (Drury *et al*., 1997; Drury *et al*., 2000; Piatti *et al*., 1995). In *Δclb2* cells, Cdc6 protein levels were suppressed throughout S-phase and particularly in mitosis (Figure 1D, 1E and 1F). This result suggests a role for Clb2 in Cdc6 stabilization during mitosis possibly by masking the Cdc6 phospho-degron. In contrast, Cdc6 was more stabilized in *cdc55-101* cells during mitosis, with a peak accumulation between 60-100 minutes (Figure 1D). Flow cytometry analysis did not show a significant difference in the cell cycle profile between wild type and *Δclb2* cells (Figure 1F). In *cdc55-101* cells, we observed a 20-minute S-phase delay, indicating that PP2A^Cdc55^ positively regulates S-phase progression (Figure 1F). We also conclude that the early Cdc6 accumulation in *cdc55-101* cells is not due to the cell cycle delay.

To examine how PP2A^Cdc55^ impacts Cdc6 localization and protein level, we analyzed Cdc6-Citrine by time-lapse microscopy in wild type and *Δcdc55* cells. Whi5 is a Start transcriptional repressor imported into the nucleus in M/G1 and exported in late G1. Whi5 nuclear import was used as a cell cycle indicator that was set as time zero, making the Start point (Figure 1G) (Doncic *et al*., 2011). Cdc6-Citrine began accumulating in the nucleus after Whi5 nuclear import and reached maximum accumulation at 9 minutes after Whi5 nuclear import in wild type cells (Figure 1H). In *Δcdc55* and *cdc55-101* cells, we observed more Cdc6-Citrine accumulation (Figure 1H). The timing of Cdc6-Citrine expression and degradation remained unchanged, as it started dropping around 15 minutes after Whi5 nuclear import in all strains (Figure 1H). This data suggests that PP2A^Cdc55^ negatively regulates Cdc6 stability in the nucleus during mitosis. We also observed Cdc6-GFP nuclear localization in early mitosis in asynchronous *cdc55-101* cells (Figure 1I, arrow). The similar Cdc6 protein accumulation patterns observed in *Δcdc55* and *cdc55-101* supports the idea that Cdc6 is solely regulated by nuclear PP2A^Cdc55^ function.

### Clb2 stabilizes Cdc6 through its N-terminus

Clb2 binds to the Cdc6 N-terminus through a conserved LxF motif (Örd *et al*., 2019). We tested if Cdc6 suppression in *Δclb2* cells is mediated through the Cdc6 N-terminus. The *CDC6-T368A* mutant contains an alanine substitution in the C-terminal phospho-degron, stabilizing Cdc6 in mitosis (Figure 2A, lanes 4+5 and 2B) (Perkins *et al*., 2001). This mutant retains the N-terminal Cdc6 phospho-degron at T39-S43. Cdc6-T368A was unstable in *Δclb2* log phase cells (Figure 2A lanes 2+3) and was more prominent in mitotic-arrested cells, supporting the idea that Clb2 shields the Cdc6 N-terminal phospho-degron (Figure 2A, lanes 5+6 and 2B) (Örd *et al*., 2019). The *CDC6-T39A-T368A* double mutant, containing mutations at both the N- and C-terminal phospho-degron, was synthetically lethal in *Δclb2* cells (Figure 2C). When Cdc6 is overexpressed and controlled under *GAL* promoter, we observed stabilized Cdc6-T39A-T368A in *Δclb2* cells during mitosis, suggesting that stabilized Cdc6 is lethal when Clb2’s inhibitory role in origin licensing is abolished (Figure S1). In contrast, GAL-Cdc6 and GAL-Cdc6-T368A were degraded in *Δclb2* cells, which suggest the importance of the Cdc6 N-terminal phospho-degron in its degradation when Clb2 is abolished (Figure S1).

**Figure 2.**
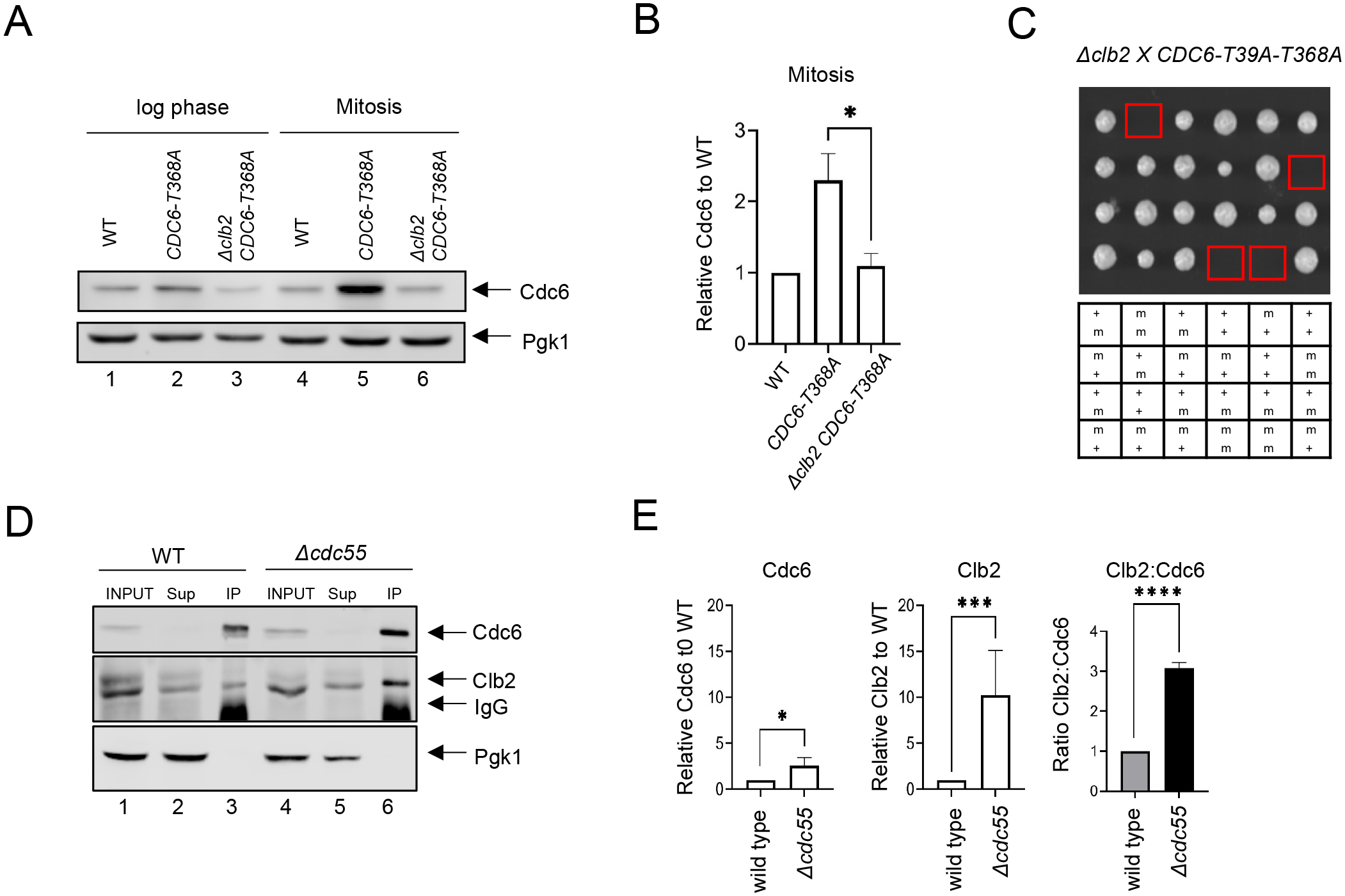
Cdc6 and Clb2 are more tightly bound in *Dcdc55* cells. A) *CDC6-PrA* (WT), *CDC6-T368A-PrA or Δclb2 CDC6-T368A-PrA* cells were incubated to log-phase or arrested in mitosis by nocodazole. Protein extracts were subjected to western blot analysis to visualize Cdc6-PrA. Pgk1 was used as a loading control. B) Cdc6 protein levels in A were quantified and normalized to WT. Three independent biological replicates were performed. An average of the relative Cdc6 band intensity is shown. Error bar is SEM (n=3; *Δclb2 CDC6-T368A-PrA* *p=0.0442 was calculated using unpaired Student’s t-test). C) *Δclb2* strain was crossed with *CDC6-T39A-T368A and* tetrad analysis was performed. The genotype for each haploid progeny is shown. + is wild type and m is mutant. Top letter is for *Δclb2* and bottom letter is for *CDC6-T39A-T368A.* D) *CDC6-9MYC* (WT) or *CDC6-9MYC Δcdc55* (*Δcdc55*) cells were arrested in mitosis by nocodazole. Samples were collected and protein was extracted (INPUT). Cdc6-9MYC was pulled down by anti-MYC agarose beads. Supernatant (Sup) or pull-down samples (IP) were analyzed by western blot to visualize Cdc6-9MYC and Clb2. Pgk1 was used as a loading control. E) Band intensities from E were quantified. Cdc6 and Clb2 levels in *Δcdc55* were normalized to those in WT. Three independent biological replicates of the experiment in E were performed. An average of the relative Cdc6 and Clb2 band intensities are shown. The ratio between Clb2 to Cdc6 was obtained and normalized to WT. Error bar is SEM (n=3; Cdc6 *Δcdc55* *p=0.0205, Clb2 *Δcdc55* ***p=0.0003, Clb2:Cdc6 *Dcdc55* ****p=0.0001 was calculated using unpaired Student’s t-test).

### Clb2-Cdc6 interaction is enhanced in *Δcdc55* cells

Cdc6 stabilization in *Δcdc55* cells indicates that PP2A^Cdc55^ inhibits Cdc6 protein accumulation. Therefore, we excluded the possibility that PP2A^Cdc55^ targets neither Cdc6 phospho-degron. We hypothesized that PP2A^Cdc55^ releases Clb2 from Cdc6, exposing the Cdc6 phospho-degrons, which subsequently leads to Cdc6 degradation. To test this possibility, we compared the Clb2:Cdc6 protein binding ratio between wild type and *Δcdc55* cells during mitosis by co-immunoprecipitation (co-IP). Cdc6 was stabilized in *Δcdc55* cells, confirming the results in Figure 1B (Figure 2D, lanes 3+6 and 2E). There was more Clb2 bound to Cdc6 in *Δcdc55* cells compared to wild type cells (Figures 2D, lanes 3+6). The relative ratio between Clb2 to Cdc6 was reproducibly higher in *Δcdc55* cells, suggesting that PP2A^Cdc55^ disrupts Clb2-Cdc6 binding (Figure 2E). From this result, we conclude that Clb2 binds tighter to a fraction of Cdc6 in *Δcdc55* cells, which may mask the Cdc6 N-terminal phospho-degron, and thereby contribute to Cdc6 stability in *Δcdc55* cells.

### PP2A^Cdc55^ targets the Cdc6 phosphorylation site T7

Our finding supports a scenario in which PP2A^Cdc55^ dephosphorylates Cdc6 at the N-terminus and disrupts Clb2-Cdc6 binding. We hypothesized that PP2A^Cdc55^ dephosphorylates Cdc6 at residue T7, which removes the Cks1 docking site, thus inhibiting Clb2 binding. We generated two *CDC6-9MYC* constructs that contain alanine mutations at the Cdk1-phosphorylation sites to test if Cdc6-T7 site is dephosphorylated by PP2A^Cdc55^. The *CDC6-6A* mutant strain contains six alanine mutations at T7A, T23A, T39A, S43A, T368A, S372A, serving as a negative control. The *CDC6-T7* mutant strain only retains the T7-phosphorylation site while all other Cdk1 phosphorylation sites are mutated to alanine (T23A, T39A, S43A, T368A, S372A) (Figure 3A). Thus, both *CDC6* constructs eliminate the function of both phospho-degrons. We raised a custom Cdc6 T7 phospho-specific antibody to detect T7 phosphorylation (Cdc6-T7p). Cdc6-T7 phosphorylation was not detected in the *CDC6-6A* mutant, confirming the specificity of the custom-made Cdc6-T7 phospho-specific antibody (Figure 3B). The Cdc6-6A protein level did not differ between *CDC55* and *Δcdc55* cells (Figure 3B). Cdc6-T7 phosphorylation was detected in the Cdc6-T7 construct which was enhanced in *Δcdc55* cells, supporting the idea that PP2A^Cdc55^ dephosphorylates the Cdc6-T7 site (Figure 3B and 3C). Cdc6-6A did not interact with Clb2, supporting a role for Cdc6 phosphorylation in protein binding (Figure 3B). Cdc6-T7 associated with Clb2 supporting our previous results that Cks1 docking to T7 is crucial for the Cdc6-Clb2 interaction (Figure 3B). The Cdc6-T7 and Clb2 interaction was enhanced in *Δcdc55* cells (Figure 3B and 3C). Next, we examined Cdc6-T7 phosphorylation during the cell cycle by alpha-factor block and release. Cdc6 was expressed during mitosis at 80 minutes in wild type and was more stabilized in *cdc55-101* (Figure 3D and 1D). We observed weak Cdc6 T7 phosphorylation in wild type cells throughout the cell cycle. In contrast, there was a spike of T7 phosphorylation in *cdc55-101* mutant cells at 80 minutes during mitosis (Figure 3D). The Cdc6 T7 phosphorylation peak coincides with Cdc6 stabilization (Figure 3D and 3E). From these results, we conclude that Cdc6 T7 phosphorylation is reversed by PP2A^Cdc55^, which releases Clb2.

**Figure 3.**
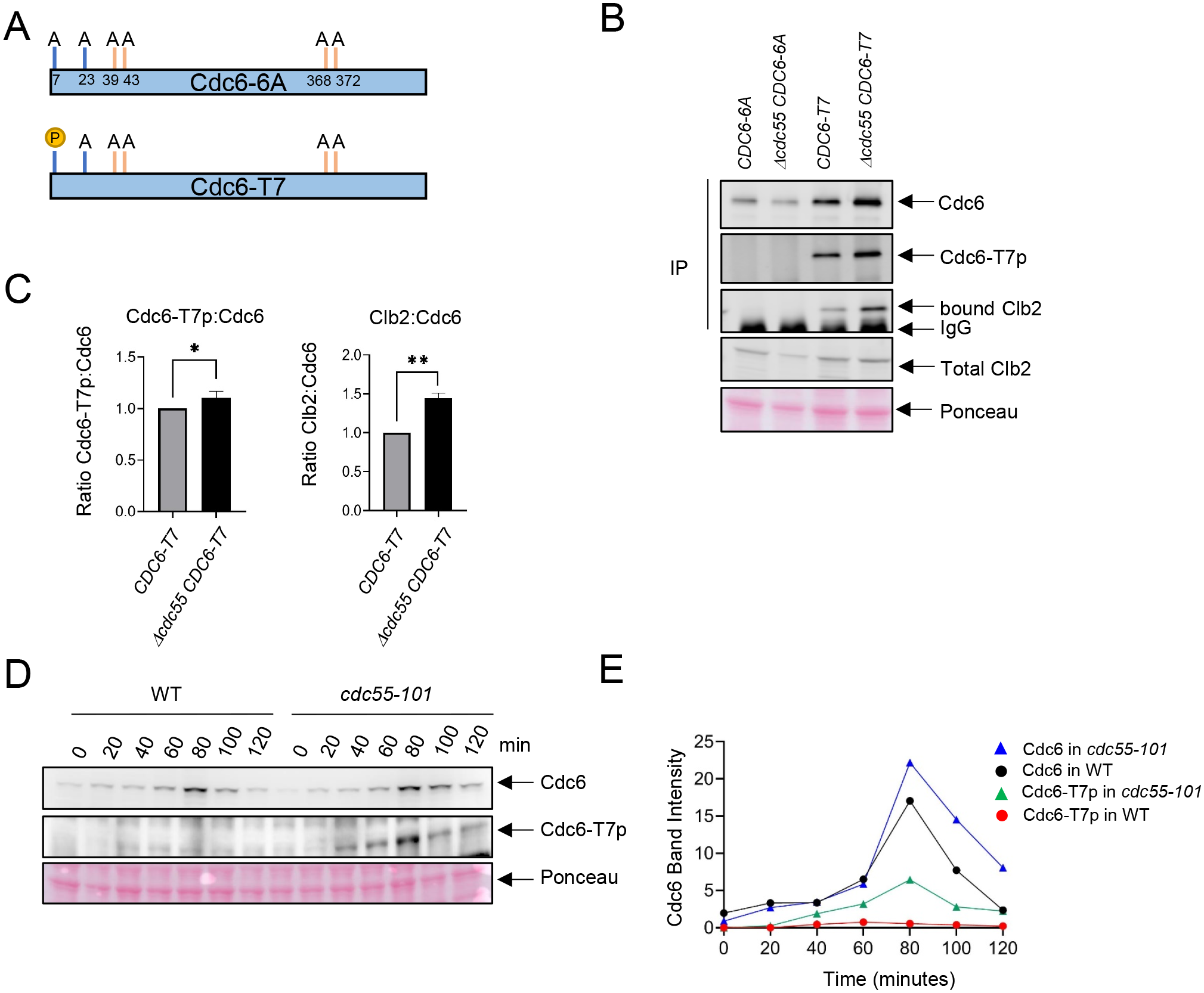
Cdc6-T7 is hyperphosphorylated in *Δcdc55* cells. A) A schematic of Cdc6 phosphorylation mutants are shown. Alanine mutations are indicated as A with amino acid positions. P indicates CDK sites accessible for phosphorylation. B) The indicated Cdc6 mutants were immuno-precipitated using anti-MYC agarose beads. A total Cdc6 and Cdc6-T7 phosphorylation were analyzed by western blot with anti-MYC and Cdc6-T7p antibodies, respectively. Clb2 was detected by anti-Clb2. C) Bands from B were quantified and the ratio between Cdc6-T7p:Cdc6 and Clb2:Cdc6 are shown. Error bars are SEM (n=3; Cdc6-T7p:Cdc6 *p=0.0468, Clb2:Cdc6 **p=0.0034 was calculated using unpaired Student’s t-test). D) *CDC6-9MYC* (*WT*) or *CDC6-9MYC cdc55-101* (*cdc55-101*) cells were arrested in G1 by alpha-factor and released. Cdc6 was visualized by anti-MYC antibody. Cdc6-T7 phosphorylation was detected by T7p phospho-specific antibody. E) Band intensities from D were quantified.

### Cdc14 stabilizes Cdc6 in late mitosis

It has been reported that Cdc14 targets Cdc6 and other pre-RC proteins (Zhai *et al*., 2010). Consequently, we examined Cdc6 protein levels in *cdc14-3* mutant cells. Cdc14 was inactivated at the non-permissive temperature upon α-factor release. We show that Cdc6 accumulation was consistently lower in *cdc14-3* compared to *cdc15-1*, an alternative mitotic mutant (Figure S2A). This result was especially prominent during mitosis at 120-180 minutes. The cell cycle profiles between *cdc14-3* and *cdc15-1* were the same, confirming that the observed difference in Cdc6 protein levels was not caused by cell cycle progression defects (Figure S2B). From these data, we conclude that Cdc14 stabilizes Cdc6 during late mitosis.

### PP2A^Cdc55^ and Cdc14 dephosphorylate Cdc6 at distinct sites

PP2A^Cdc55^ and Cdc14 exhibited opposite effects on Cdc6 stability (Figure 1 and Figure S2). We hypothesized that Cdc14 may target a Cdc6 phospho-degron, leading to Cdc6 stabilization. PP2A^Cdc55^ and Cdc14 specificity towards Cdc6 was examined by *in vitro* phosphatase assays. Recombinant Cdc6 was preincubated with Clb5-Cdk1 in the presence of [γ-^32^P]-ATP. The reaction mixture was further incubated with purified Cdc14 or PP2A^Cdc55^ for dephosphorylation. Purified Sic1 was also added to the mixture in order to inhibit further phosphorylation of Cdc6 by Clb5-Cdk1 after addition of the phosphatases. The intensity of [^32^P]-label on Cdc6 was measured by autoradiography. Full-length wild type Cdc6 was moderately dephosphorylated by both PP2A^Cdc55^ and Cdc14 (Figure 4A), indicating that Cdc6 is directly targeted by these phosphatases. The phosphate on Cdc6-T7 was efficiently removed by PP2A^Cdc55^ when T7 was the only phosphorylation site available, but not by Cdc14 (Figure 4B). The same trend was observed with Cdc6-T23, but to a lesser extent (Figure 4C). When the Cdc6 C-terminal phospho-degron was the only phosphorylation site available, it was dephosphorylated by Cdc14, but not by PP2A^Cdc55^ (Figure 4D). N-Swe1 and Csa1 were used as positive controls to show PP2A^Cdc55^ (Harvey *et al*., 2011) and Cdc14 specificity (Örd *et al*., 2020), respectively (Figure 4E and 4F). Neither Cdc6-T39 nor -S43 was dephosphorylated by PP2A^Cdc55^ or Cdc14, indicating that the Cdc6 N-terminal phospho-degron is not controlled by these phosphatases (Figure S3). These results suggest that PP2A^Cdc55^ directly dephosphorylates Cdc6-T7 and -T23, while Cdc14 dephosphorylates the Cdc6 C-terminal phospho-degron at T368-S372. Thus, PP2A^Cdc55^ and Cdc14 showed sequence specificity towards Cdc6.

**Figure 4.**
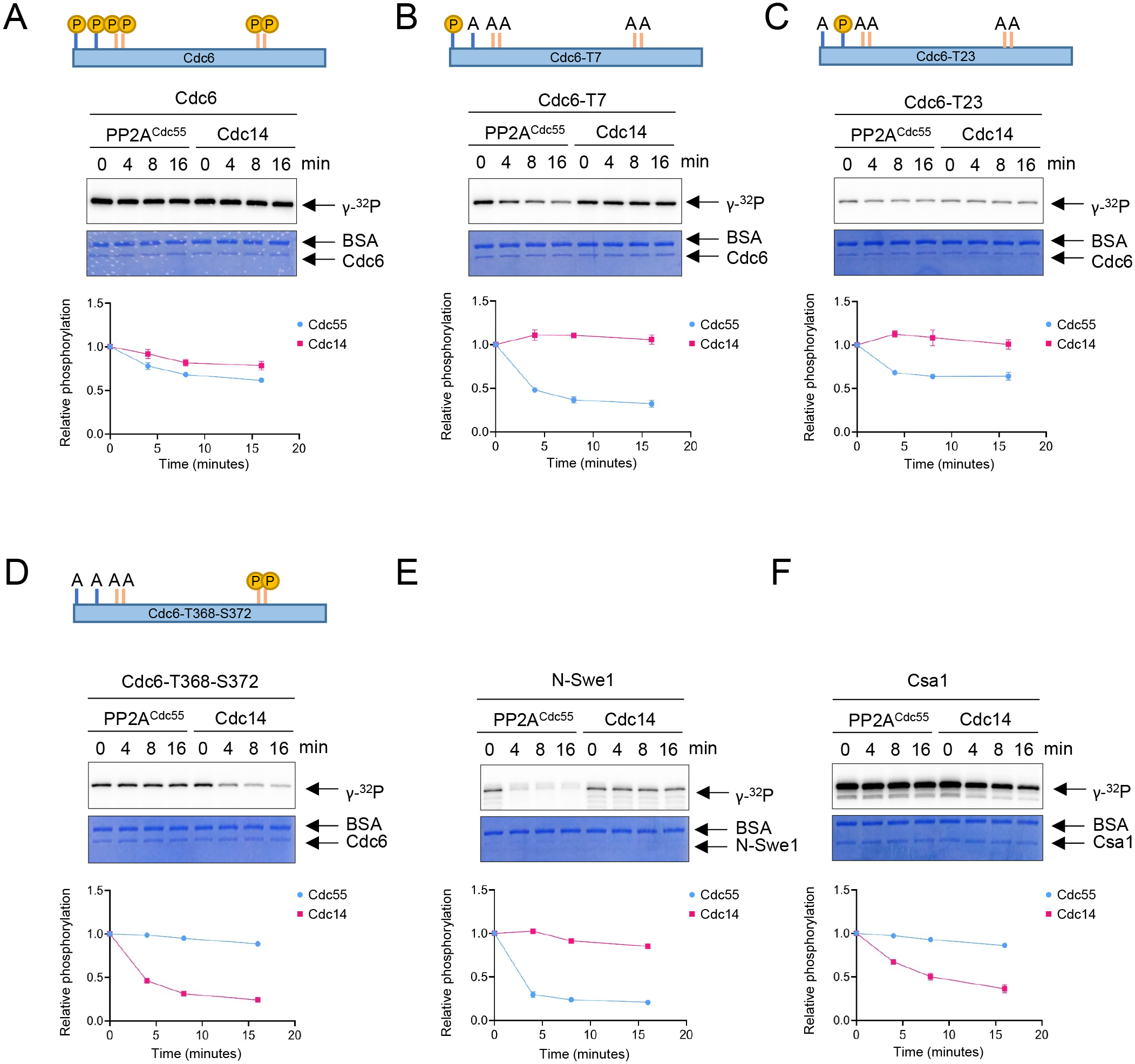
PP2A and Cdc14 dephosphorylate Cdc6-T7, –T23 and T368-S372, respectively. Full length Cdc6, Cdc6-T23A-T39A-S43A-T368A-S372A (Cdc6-T7), Cdc6-T7A-T39A-S43A-T368A-S372A (Cdc6-T23), Cdc6-T7A-T23A-T39A-S43A (Cdc6-T368-S372) were phosphorylated for 60 minutes at room temperature using purified Clb5-Cdk1 in the presence of [γ-^32^P]-ATP. At time 0, Sic1 and phosphatase Cdc14 or PP2A^Cdc55^ were added to the reactions to inhibit Clb5-Cdk1. A time course was taken, and samples were measured for dephosphorylation. N-Swe1 was used as a positive control for PP2A^Cdc55^ and Csa1 for Cdc14. The samples were loaded on SDS-PAGE and following electrophoresis, the gels were stained using Coomassie Brilliant Blue. Signals were quantified using ImageQuant TL. Three independent biological replicates of the phosphatase assays were performed. Error bar is SEM (n=3).

### Sic1 releases Clb2 from Cdc6

Deletion of *CDC55* slows down S-phase progression. However, cells were still capable of completing S-phase (Figure 1F). The key regulator of Clb2-Cdk1 inhibition, Sic1, is a promoter of origin licensing in G1 in conjunction with APC^Cdh1^ (Lengronne & Schwob, 2002; Wäsch & Cross, 2002). It is possible that Sic1 also drives origin licensing through Cdc6. Both Cdh1 and Sic1 act in overlapping mechanisms that promote mitotic exit that is potentially coupled to origin licensing (Visintin *et al*., 1998). We hypothesized that this is due to high level of Clb2 binding to Cdc6. A *CDC6-T39A-T368A Δcdh1* strain was used to obtain stable Cdc6 expression. Transient Sic1 expression was induced in a *GAL-SIC1* strain using galactose. Overexpression of Sic1 released Clb2 from Cdc6 in mitotic cells arrested by nocodazole (Figure 5A and 5B).

**Figure 5.**
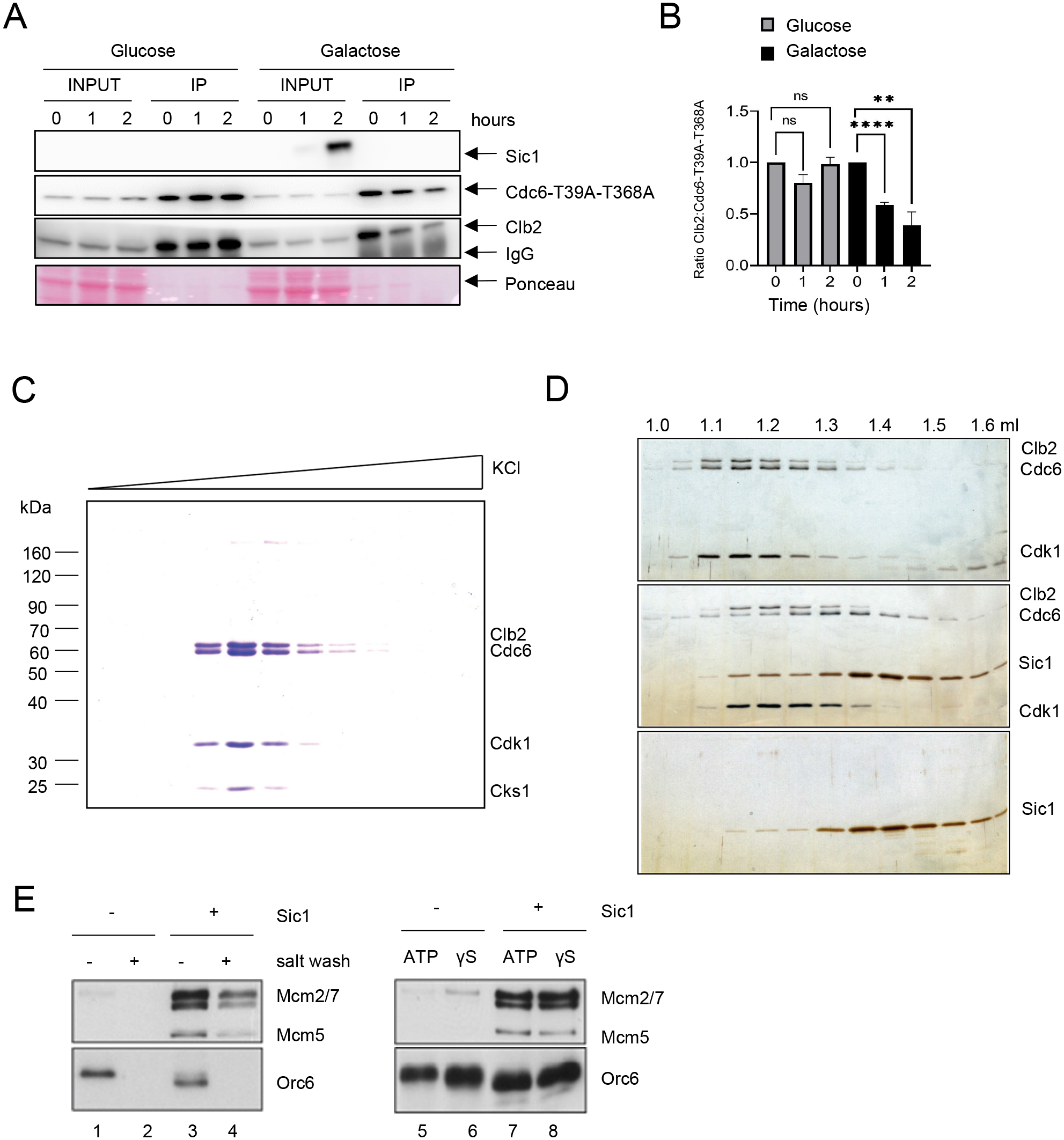
Sic1 promotes origin licensing by releasing Clb2/CDK from Cdc6. A) *GAL-SIC1-HA CDC6-T39A-T368A-PrA Δcdh1* cells were arrested in mitosis by nocodazole for 2 hours before the addition of glucose or galactose. Samples were collected at the indicated time points and protein extracted (INPUT). Cdc6-T39A-T368A-PrA pulled down by anti-IgG Dynabeads was analyzed by western blot to visualize Sic1, Cdc6-T39A-T368A, and Clb2. B) Clb2:Cdc6-T39A-T368A IP ratio relative to time 0 from A is shown. Error bar is SEM (n=3; Glucose p=0.8414, p=0.0729, ns, not significant; Galactose ****p<0.0001, **p= 0.0090, was calculated using unpaired Student t-test). C) Cdc6 was expressed and purified from M-phase cell extracts. The elution profile of the Clb2·Cdk1·Cks1·Cdc6 complex after ion-exchange chromatography is shown. D) Purified Clb2·CDK·Cdc6 complex and Sic1 were analyzed by gel-filtration. Fractions were analyzed by SDS-PAGE and silver staining. Elution positions are indicated on the top. Gel-positions of indicated proteins are located on the right. E) MCM loading assays with purified ORC, Clb2·Cdk1·Cks1·Cdc6, Cdt1·Mcm2-7, and Sic1 on origin DNA immobilized on paramagnetic beads. Loading was performed in the presence or absence of Sic1 and the presence of either a salt wash (left) or ATP and ATP analog, ATPγS (γS) (right).

Next, we tested if Sic1 promotes Mcm2-7 DNA loading using a biochemical approach with purified proteins (Ayuda-Durán *et al*., 2014; Remus *et al*., 2009; Seki & Diffley, 2000). The stringent regulation of intracellular Cdc6 protein levels by proteolytic degradation in both G1 and S phase severely hampers the purification of endogenous Cdc6 from these cell cycle stages even after galactose-induced overexpression. However, we observed that extended overexpression of TAP^TCP^-tagged Cdc6 upon release from α-factor arrest induces a mitotic arrest phenotype in yeast cells, characterized by a 2C DNA content and the presence of a large elongated bud (data not shown). Intriguingly, Cdc6 isolated from whole-cell extracts of such mitotically arrested cells exists in a stable stoichiometric complex with three additional proteins that we identified as Clb2, Cdk1 and Cks1 by mass-spectrometry (Figure 5C). The formation of this complex after Cdc6 overexpression is consistent with the previous observation of a Cdc6·Clb2·Cdk1 complex in G2/M cells (Mimura *et al*., 2004). Importantly, using gel-filtration analysis, we find that Cdc6 is released from the Clb2·Cdk1·Cks1 complex in the presence of the CDK inhibitor Sic1, which itself forms a stable complex with Clb2·Cdk1·Cks1 (Figure 5D).

Clb2-Cdk1 binding to Cdc6 has been proposed to inhibit the origin licensing activity of Cdc6 as part of cellular re-replication control mechanisms (Mimura *et al*., 2004). We tested the ability of Cdc6·Clb2·Cdk1·Cks1 to promote Mcm2-7 loading onto DNA in conjunction with purified ORC and Cdt1·Mcm2-7, analogous to experiments described previously with purified Cdc6 (Remus *et al*., 2009). Using this approach, we find that Cdc6·Clb2·Cdk1·Cks1 does not support the formation of salt-stable MCM complexes on DNA, indicating that Cdc6 in complex with Clb2·Cdk1·Cks1 is inactive for Mcm2-7 loading (Figure 5E, lanes 1+2). Importantly, normal Mcm2-7 loading was observed in reactions containing both purified Sic1 and Cdc6·Clb2·Cdk1·Cks1 (Figure 5E, lanes 3+4), indicating that Clb2·Cdk1·Cks1 sequesters Cdc6 and that this form of Cdc6 inhibition can be reversed by release of Cdc6 from the Cdc6·Clb2·Cdk1·Cks1 complex by Sic1. However, we noted that Orc6 is efficiently phosphorylated in the presence of Cdc6·Clb2·Cdk1·Cks1, as indicated by the pronounced gel-mobility shift of Orc6 (Figure 5E, compare lanes 1+3). This Orc6 phosphorylation is inhibited in the presence of Sic1, indicating that Clb2-Cdk1 retains partial kinase activity in the Cdc6·Clb2·Cdk1·Cks1 complex. It is known that CDK phosphorylation inhibits the Mcm2-7 loading function of ORC (Frigola *et al*., 2013). It was possible that the inhibition of Mcm2-7 loading in the presence of Cdc6·Clb2·Cdk1·Cks1 was due to ORC phosphorylation and not Cdc6 sequestration. However, we noted that not only Mcm2-7 loading, but even Mcm2-7 recruitment to ORC/Cdc6, which can be monitored in the presence of ATPγS, is inhibited in the presence of Cdc6·Clb2·Cdk1·Cks1 (Figure 5E, lanes 5+6), and this inhibition is again reversed in the presence of Sic1 (Figure 5E, lanes 7+8). Since ORC phosphorylation specifically inhibits Mcm2-7 loading but not Mcm2-7 recruitment (Frigola *et al*., 2013), this data demonstrates that the mechanism of licensing inhibition by Cdc6·Clb2·Cdk1·Cks1 complex formation is distinct from that by CDK phosphorylation of ORC. In summary, we conclude that Clb2-Cdk1 inhibits Cdc6 by physical sequestration and this inhibition is reversible by the Sic1-mediated release of Cdc6 from the Cdc6·Clb2·Cdk1·Cks1 complex.

### Key functional residues/motifs of Cdc6 and Clb2 are located within regions of intrinsic disorder

The necessary structural elements in Cdc6 responsible for protein binding are not well understood. Protein binding motifs often fall in intrinsically <u>d</u>isordered regions (IDRs), which are not well characterized structurally due to the limitations of crystal structure analysis (Fuxreiter *et al*., 2007; Wright & Dyson, 2015). IDRs in Cdc6 were identified by several programs including IUPred (Mészáros *et al*., 2018) and Anchor (Mészáros *et al*., 2009), which include: 1) Cdc6-N terminus residues 1-59, and 2) Cdc6 C-terminus residues 350-378 (Figure 6A and S4A). Next, we constructed a full-length Cdc6 structure model by extracting the known structure from Protein Data Bank (PDB ID: 5V8F) (Yuan *et al*., 2017), which was lacking some of the key functional residues and regions e.g., 1-59, 68-80, 129-163 and 349-387. Indeed, these regions fall into the IDRs identified in Figure 6A. We generated a complete structural representation of Cdc6 through predictions of both IDRs and secondary structure combined with template-based modelling and *ab initio* approaches (See Methods). The missing regions modeled in the Cdc6 N-terminus span residues 1-59 (Figure 6B, left) as well as the centrally located residues 350-378 (Figure 6B, right), which correspond to the predicted IDRs (Figure 6A). The Cdc6 N-terminal IDR comprises four Cdk1 consensus sites (T7, T23, T39, S43) and two cyclin binding motifs RxL (29-31) and LxF (47-49), all of which are critical for Cdc6 phospho-regulation (Figure 6B, left). The Cdc6 C-terminal IDR (residues 350-378) contains phospho-degron 368-372 (Figure 6B, right). We also compared our Cdc6 structural model with the recently released AlphaFold and evaluated the model quality using Verify 3D software (Figure S4B). The 3D-1D profiles from Verify 3D measure the compatibility of the primary structure of a protein with its three-dimensional structural environment in the model. An average 3D-1D score of 0.2 (shown by the green line in the plots) denotes the threshold of acceptable model quality i.e., scores around this threshold or above are inferred as correctly modeled regions (Figure S4C). The phospho-degrons (red boxes) scored close to or at the 0.2 threshold in our model, while both regions score well below the threshold in the Alphafold model (Figure S4C). This is also consistent with low confidence scores (less than 70) generated by AlphaFold’s internal confidence rating (Figure S4B, yellow and orange). The scores in the structured regions of both models are comparable. Thus, our model outperforms the AlphaFold model in the region of the modeled IDRs (Figure S4C).

**Figure 6.**
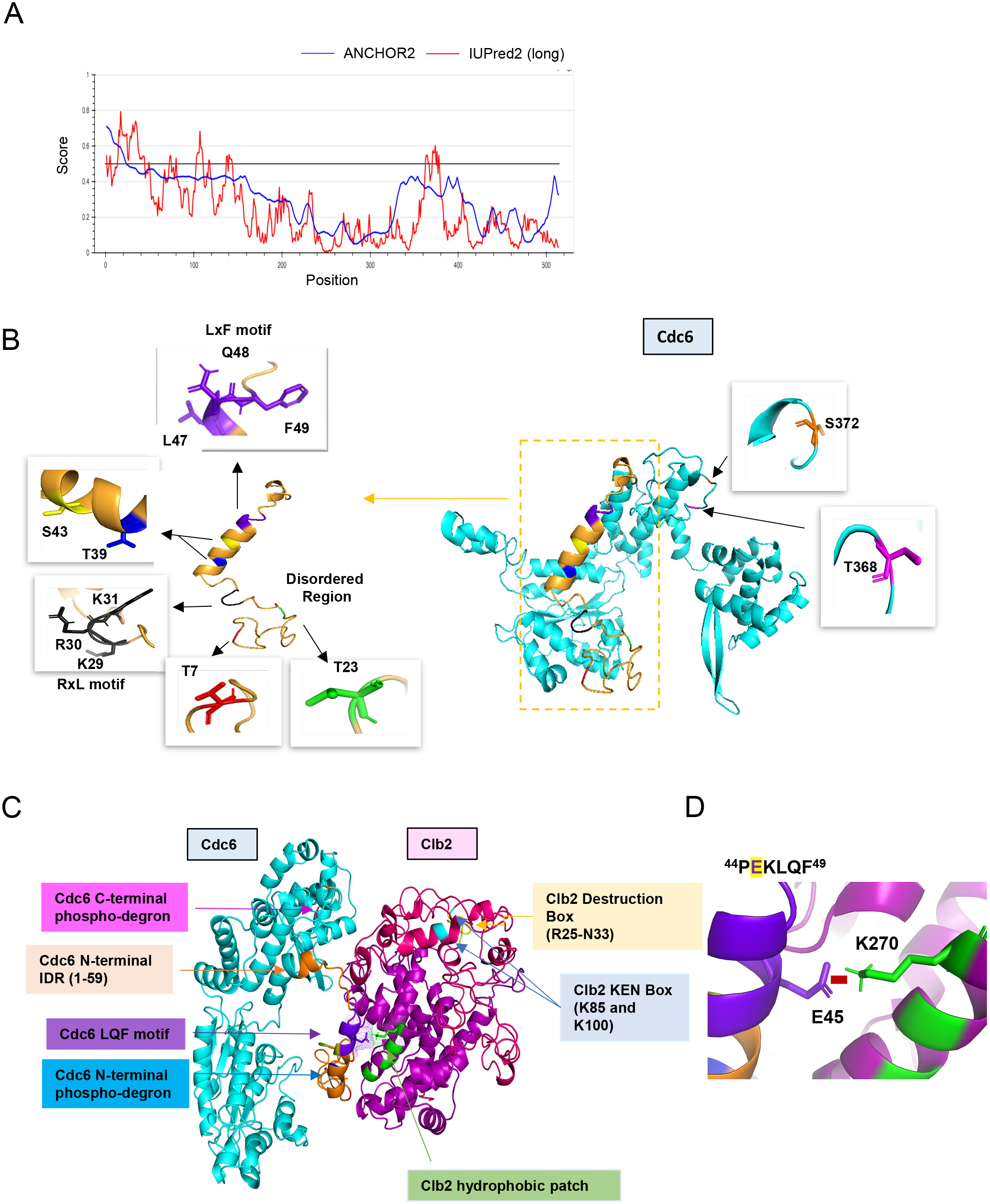
Predicted Cdc6-Clb2 structure by bioinformatics. A) Disordered regions were mapped using ANCHOR2 and IUPred2 software. B) A full length Cdc6 protein structure composed of 513 amino acids was predicted based on the amino-acid sequence and known structure information (cyan and orange on the right). N-terminal highly disordered region spans in 1-59 residues (orange box and highlighted on the left). T7 is in red, T23 is in green, RxL motif is in black and LxF motif in purple. N-terminal phospho-degron is located at T39-S43 and C-terminal phospho-degron is at T368-S372. C) Cdc6-Clb2 binding structure with Cdc6 IDR (orange), Cdc6 LQF motif (purple), Cdc6 N-terminal phospho-degron (blue), Cdc6 C-terminal phospho-degron (pink), Clb2 D-box (yellow), Clb2 KEN Box (light blue), and Clb2 hydrophobic patch (green). D) Cdc6–Clb2 interaction through Cdc6 E45 (purple) and Clb2 K270 (green). Red bar shows salt bridge.

Finally, we predicted a potential scenario of Cdc6-Clb2 interaction, where Cdc6 E45 interacts with Clb2 K270 in hydrophobic patch via a salt bridge (Figure 6C and 6D) that highlights the importance of the IDR in this interaction. We speculate that a transiently structured helix adopted by the PEKLQF motif in our Cdc6 model may be important for its interaction with the hydrophobic patch in Clb2. The N-terminus of Clb2 located from residues 1-210 (data not shown) contained an IDR which includes the KEN box (residue 85 and 100) and the destruction box (D-box) (residues 25-33), which are both necessary for Clb2 degradation via APC (Hendrickson *et al*., 2001; Meyn Iii *et al*., 2002) (Figure 6C). The ordered regions of Clb2 include the hydrophobic patch (N260, L264, Y267, K270) that facilitates protein binding to its substrate such as Cdc6 (Hunt, 1991; Örd *et al*., 2019) (Figure 6C). Importantly, our Cdc6-Clb2 protein interaction model also captured Cdc6 N-terminal phospho-degron protected by Clb2, thereby inhibiting Cdc6 degradation.

## Discussion

### Cdc6 regulation by PP2A^Cdc55^

In this study, we showed that PP2A^Cdc55^ dephosphorylates Cdc6 at T7 and T23 to release Cdc6 from Clb2·Cdk1·Cks1 complex. Neither PP2A^Cdc55^ nor Cdc14 dephosphorylated the Cdc6 N-terminal phospho-degron, suggesting that there are additional mechanisms involved (Figure S3). The Cdc6 T7 site is located in an IDR, where PP2A^Cdc55^ and Cks1 might gain easier access for docking (Figure 6A and 6B), which may trigger a cascade of Cdc6 dephosphorylation in early mitosis (Touati *et al*., 2019). Our study also supports a model in which PP2A^Cdc55^ prefers phospho-threonines over phospho-serines (Godfrey *et al*., 2017).

Previous work in *Xenopus* egg extracts indicated that immunodepletion of the trimeric PP2A complex inhibits replication initiation without regulating pre-RC assembly (X.-H. Lin *et al*., 1998). A follow-up study suggested that PP2A targets proteins required for DNA replication initiation but not the elongation and termination proteins such as Cdc45 (Chou *et al*., 2002). Sld2 and Sld3 are potential substrates for PP2A (Bloom & Cross, 2007). Therefore, PP2A might regulate multiple replication proteins at different steps. Mammalian PP2A is regulated by a diverse group of regulatory subunits, termed, B, B’, B’’ and B’’’ (Janssens & Goris, 2001). B55 is a mammalian homologue of Cdc55 which plays a role in mitosis (Schmitz *et al*., 2010). PP2A^B55^ also selectively targets phospho-threonines over phospho-serines *in vitro* (Agostinis *et al*., 1990; Cundell *et al*., 2016). PR48 (B’’ family) has been identified as a Cdc6 binding partner through yeast two-hybrid screening (Yan *et al*., 2000). PR48 mediates protein interaction with the Cdk1 phosphorylation site-containing Cdc6 N-terminus (Yan *et al*., 2000). PR70, another B’’ family subunit, has also been implicated in Cdc6 regulation by enhancing phosphatase activity towards Cdc6 (Davis *et al*., 2008; Wlodarchak *et al*., 2013; Yan *et al*., 2000). Thus, our data define Cdc6 dephosphorylation by PP2A in a simple system, which might be a conserved mechanism in eukaryotes.

### Cdc14 and DNA replication

The Cdc6 C-terminal phospho-degron is targeted by SCF^Cdc4^ for protein degradation in mitosis (Drury *et al*., 2000). Here, we show that the Cdc6 C-terminal phospho-degron is dephosphorylated by Cdc14 to stabilize Cdc6 (Figure 4). Cdc14 is inhibited by PP2A^Cdc55^ through Net1 dephosphorylation (Queralt *et al*., 2006), which indicates that Cdc14-dependent Cdc6 dephosphorylation is not triggered until Clb2-Cdk1 activity overrides PP2A^Cdc55^ activity. An elevated activity of Clb2-Cdk1 phosphorylates Net1 to release Cdc14 nucleolus (Figure 7B). PP2A^Cdc55^-dependent Cdc6 T7 dephosphorylation releases Clb2 which may enhance Net1 phosphorylation because Clb2 is free from its inhibitor Cdc6 (Figure 7B left). Once Cdc14 is released from Net1, Cdc14 dephosphorylates Cdc6, Cdh1 and Sic1 to license the origin (Figure 7B right). This temporal regulation of Cdc14 ensure that Cdk1 substrates are sequentially dephosphorylated (Kataria *et al*., 2018). Such a temporal regulation is also achieved by cell cycle-dependent Cdc55 localization; nuclear Cdc55 inhibits metaphase-anaphase transition through APC inhibition (Queralt & Uhlmann, 2008; Rossio *et al*., 2013).

**Figure 7.**
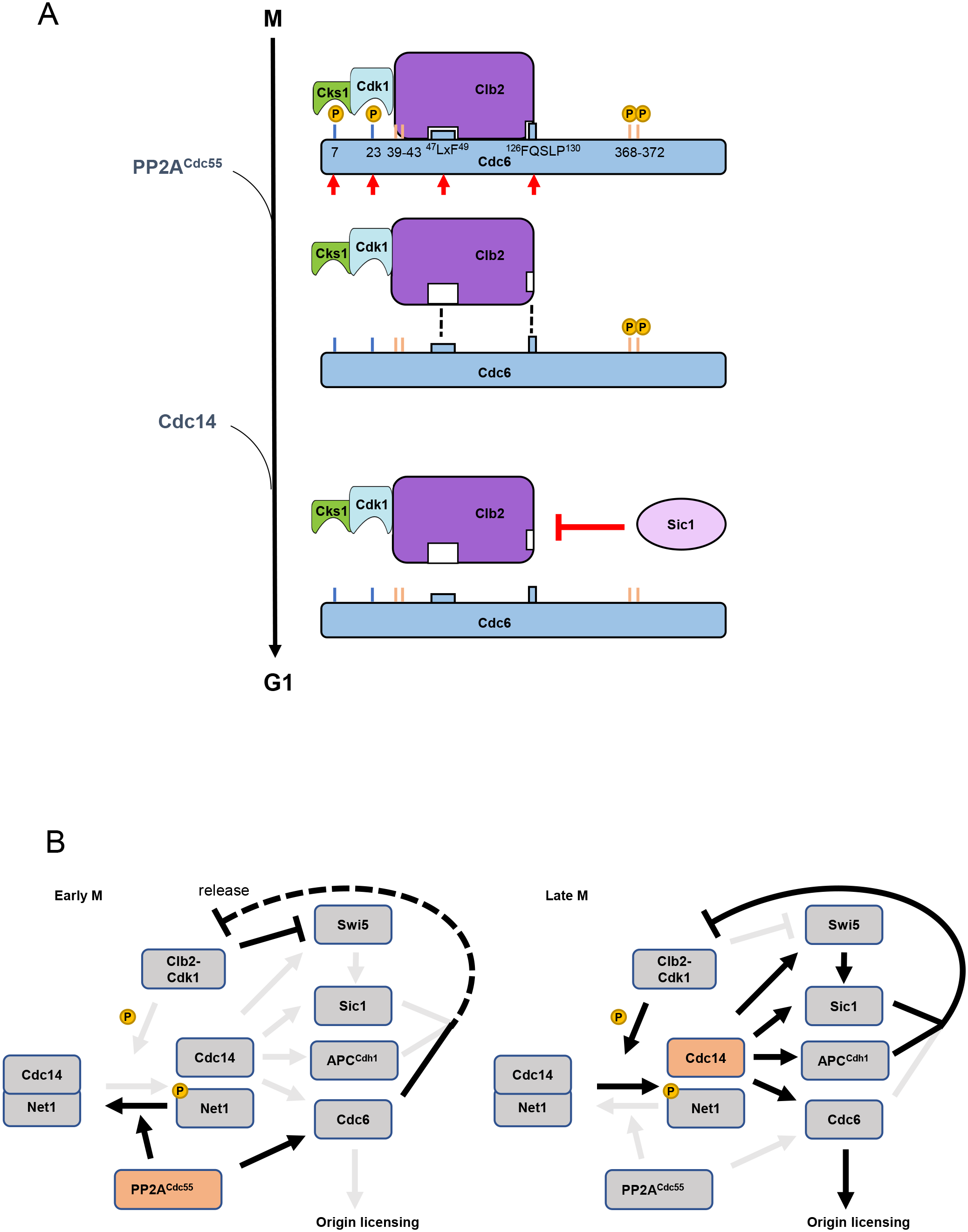
A model of Cdc6 regulation by Clb2, PP2A^Cdc55^, Sic1, and Cdc14. A) In mitosis, the Clb2·Cdk1·Cks1 complex binds to phosphorylated Cdc6 N-terminus to mask the phospho-degron and prohibit Cdc6 loading on DNA, which is mediated by the LxF motif and FQSLP enhancing region. PP2A^Cdc55^ dephosphorylates Cdc6-T7 and – T23 sites to disrupt Clb2·Cdk1·Cks1·Cdc6 complex. Sic1 releases Clb2·Cdk1·Cks1 from Cdc6 to promote origin licensing. Cdc14 dephosphorylates Cdc6 C-terminal phospho-degron for stabilization. Cks1 docking sites T7 and T23 are colored blue whereas phospho-degrons are colored orange. Red arrows show sites and motifs where Clb2-Cdk1 inhibits Cdc6 loading. Dashed lines indicate reduced binding. B) In early mitosis, PP2A^Cdc55^ inhibits Cdc14 release and promotes a rise in Clb2-Cdk1 activity by dephosphorylating Cdc6. During late mitosis, Clb2-Cdk1 initiates Cdc14 release from inhibitory binding by Net1. Cdc14 targets Swi5, Sic1, Cdh1, and Cdc6, contributing to full suppression of Clb2-Cdk1 activity and origin licensing.

There is evidence that Cdc14 is a key regulator for promoting DNA replication. When Cdc14 is not properly sequestered in *CDC14-TAB6* mutant, cells show DNA replication defects when *CLB5* is deleted (Bloom & Cross, 2007). Cdc14 dephosphorylates replication initiation proteins Orc2, Orc6, Cdc6, and Mcm3 (Zhai *et al*., 2010). Ectopic Cdc14 expression induces DNA re-replication, supporting the idea that Cdc14-dependent dephosphorylation on pre-RC components is a requirement for origin licensing (Zhai *et al*., 2010). Thus, Cdc14 is a key component to assemble pre-RCs in late mitosis.

### Clb2 function in Cdc6

Our results show that Clb2·Cdk1·Cks1 binding to Cdc6 may hinder SCF^Cdc4^ recognition of the N-terminal phospho-degron and consequently leads to Cdc6 stabilization (Figure 2 and 6C). This indicates that the Cdc6 C-terminal phospho-degron is the key element for Cdc6 degradation until Clb2 is released. We have previously shown that the Cdc6 C-terminal phospho-degron is targeted by Mck1, a yeast homologue of GSK3, for protein degradation (Al-Zain *et al*., 2015; Ikui *et al*., 2012).

Despite high Clb2-Cdk1 activity and an exposed phospho-degron, Cdc6 can be partially stabilized in mitosis until Mck1 phosphorylates Cdc6-T368. It is of interest to study how Mck1 is activated during the cell cycle. Mck1 is involved in DNA damage signaling (Al-Zain *et al*., 2015; Li *et al*., 2019), therefore the Cdc6 C-terminus, which is available throughout the cell cycle, plays a role in controlling the timing of licensing under stress conditions.

Clb2 degradation is mediated through Anaphase Promoting Complex (APC) coupled with its activator Cdh1 through KEN- and D-boxes. Cdh1 is inhibited until it is dephosphorylated by Cdc14. KEN- and D-boxes are located in IDRs, which may also enhance interaction with APC to facilitate Clb2 degradation upon mitotic exit (Figure 6C) (Radivojac *et al*., 2010).

### Sic1 regulation

We showed that Sic1 releases Clb2 from Cdc6 which supports origin licensing (Figure 5). Sic1 contains multiple Cdk1 phosphorylation sites, which includes a phospho-threonine Cks1 docking site at the N-terminus (Kõivomägi *et al*., 2011; Venta *et al*., 2020). Cks1 docking triggers an N- to C-directed multisite phosphorylation cascade both in Sic1 and Cdc6, which results in the creation of phospho-degrons targeted by SCF^Cdc4^ (Örd *et al*., 2019; Venta *et al*., 2020). Thus, Cdc6 and Sic1 share a functional similarity in their phosphorylation patterns. Since PP2A^Cdc55^ dephosphorylates Cks1 docking site T7 on Cdc6, it is worth examining if PP2A^Cdc55^ dephosphorylates Sic1 at the N-terminal Cks1 docking site. Sic1, Cdh1 and Cdc6 are Clb2-Cdk1 inhibitors as well as Cdc14 substrates and SCF^Cdc4^ targets (Calzada *et al*., 2001; Örd *et al*., 2019; Verma *et al*., 1997; Visintin *et al*., 1998). Cdc14-dependent Swi5 dephosphorylation also activates Sic1 nuclear import and transcription at the end of mitosis (Visintin *et al*., 1998). Thus, Cdc14 release from the nucleolus targets at least four components to achieve a complete inhibition of Clb2-Cdk1 activity during mitotic exit, leading to origin licensing (Figure 7B).

### Short Linear Motifs (SLiMs) and IDRs

Recent studies have expanded the role of cyclin-specific <u>s</u>hort <u>li</u>near <u>m</u>otifs (SLiMs), which are short functional sequence motifs often found in IDRs. SLiMs such as RxL and LxF motifs dictate Cdk1 substrate specificity and control cell cycle progression throughout the cell cycle (Bhaduri & Pryciak, 2011; Faustova *et al*., 2021; Örd *et al*., 2020; Örd *et al*., 2019). This is also conserved in mammalian cells (Allan *et al*., 2020; Takeda *et al*., 2001; Topacio *et al*., 2019). Phosphatases also utilize SLiMs for substrate docking. In yeast, Cdc14 scaffolds to a PxL motif on its substrates to mediate dephosphorylation events (Kataria *et al*., 2018). Currently, there is no identified docking motif for PP2A^Cdc55^ in yeast. Mammalian homolog PP2A^B55^ recognizes a polybasic docking sequence that flanks CDK phosphorylation sites (Cundell *et al*., 2016). PP2A^B56^ specific SLiMs are defined as LSPIxE in humans (Wang *et al*., 2016). Future research should focus on docking motifs for phosphatases and their regulatory subunits to understand phosphatase specificity in yeast.

Our data illustrate multiple IDRs on Cdc6 (Figure 6A). Notably, the N-terminal Cdc6 IDR include SLiMs such as RxL and LxF motifs, which supports a model that depicts the Cdc6 N-terminus as a hub for multiple protein-protein interactions (Figure 6B). Cdc6 IDRs are readily available for protein binding in G1/S, during which Cdc6 disengages from ORC and is subjected to SCF-dependent degradation (Drury *et al*., 1997; Piatti *et al*., 1995). Cdc6 is then shifted towards an origin licensing preparation period where Cdc6 is stabilized but not licensed via Clb2 binding. This temporal mode is released when Cdc14 outcompetes with Clb2-Cdk1 activity at the end of mitosis.

In humans, Cyclin A-CDK2 was shown to utilize SLiMs to promote interaction with Cdc6 and ORC (Hossain *et al*., 2021). Our studies on Cdc6 regulation shed light on the dynamics of phosphorylation and dephosphorylation that drive DNA replication initiation upon mitotic exit. In this study, we showed first evidence to link PP2A with DNA replication. This study also provided a stepwise origin licensing process mediated by PP2A, Cdc14 and Sic1.

### Methods Yeast cultures

Yeast extract peptone medium with glucose (YPD) was used to grow cells for western blot and flow cytometry analysis. Low fluorescence medium with glucose supplemented with adenine was used to visualize Cdc6-GFP under fluorescence microscope (Sheff & Thorn, 2004). G1 arrest was achieved by α-factor at 50 nM for 2 hours at 30°C. Cells were washed three times and resuspended in YPD to release the G1 arrest. Mitotic arrest was achieved by nocodazole at 15μg/ml for 2 hours at 30°C.

### Yeast strains

Standard methods were used for mating, tetrad analysis and transformation. All yeast strains are haploid congenic to W303 background. Strain list in this study is in Supplemental Table 1.

### Western blotting, co-IP and Phos-tag assay

Cells were lysed in TBT buffer containing protease inhibitors and Phos STOP (Roche) with acid-washed glass bead agitation using Fast-Prep as previously described (Bloom *et al*., 2011). Proteins were separated by SDS-PAGE with Novex 4–20% Tris-glycine polyacrylamide gel (Life Technologies). Cdc6-9MYC was detected using anti-cMYC antibody 9E10 (Sigma-Aldrich) at 1:5000 dilution and anti-Pgk1 (Thermo Fisher) at 1:5000 as a loading control. Cdc6-protein A was detected by HRP-conjugated rabbit IgG (Sigma). Anti-Clb2 antibody is a gift from Frederick Cross. Images were developed using a Fuji LAS 4000 Imager (GE Healthcare Life Sciences, Pittsburgh, PA) or Odyssey CLx imager (LI-COR Lincoln, NE). Co-immunoprecipitation was performed with rabbit anti-MYC conjugated agarose beads (Sigma-Aldrich, St Louis, MO) for 1 h at 4°C. Cdc6-T7 phospho antibody is custom antibody raised in rabbit (Covance, Princeton, NJ). Phos-tag analysis was performed using 8% phos-tag acrylamide gels as previously described (Fujifilm Wako Pure Chemical, Osaka, Japan) (Kinoshita *et al*., 2009; Kinoshita *et al*., 2006; Link & LaBaer, 2011)

### FACS

Cells were fixed and stained with propidium iodide staining as previously described (Epstein & Cross, 1992). Flow cytometry analysis was performed using a BD Accuri C6 flow cytometer (BD Biosciences). A total of 20,000 cells were analyzed per sample.

### Protein purification for *in vitro* phosphatase assay

Cdc6, Sic1, Swe1(1-450) and Cdc14 were expressed in *E. coli* BL21RP cells from pET28a-based vectors as fusions with 6xHis tag. Csa1 was expressed as GST fusion in BL21RP cells from pGEX-4T1-based vector. The expression of Cdc6 and Swe1(1-450) was induced at 23 °C using 0.3 mM IPTG, Sic1 at 37 °C using 1 mM IPTG, Cdc14 at 23 °C using 0.125 mM IPTG, and Csa1 at 16 °C using 0.3 mM IPTG. 6xHis-Cdc6, 6xHis-Sic1 and 6xHis-Swe1(1-450) were purified by standard cobalt affinity chromatography with 200 mM imidazole used for elution. 6xHis-Cdc14 was purified using nickel affinity chromatography with 250 mM imidazole used for elution. GST-Csa1 was purified using Glutathione Sepharose (GE Healthcare). Clb5-Cdk1 complex was purified from *S. cerevisiae* culture where TAP-tagged Clb5 was overexpressed from *GAL1* promoter as described previously (Puig *et al*., 2001; Ubersax *et al*., 2003). The yeast lysate was prepared using Mixer Mill MM 400 (Retch). PP2A^Cdc55^ complex was purified from yeast cells containing 3HA-tagged Cdc55 (Anastasia *et al*., 2012). The yeast culture was grown in YPD to OD_600_=1.4, when the cells were collected and snap frozen. The cells were lysed using Mixer Mill MM 400. The lysate was cleared by centrifugation and the supernatant was incubated with anti-HA agarose beads for 3 hours. The beads were washed thoroughly and PP2A^Cdc55^ was eluted with buffer containing HA dipeptide.

### *In vitro* dephosphorylation assay

Substrate proteins Cdc6, Swe1(1-450) and Csa1 at 500 nM concentration were phosphorylated for 60 minutes at room temperature using 2 nM Clb5-Cdk1 in a buffer containing 50 mM HEPES-KOH, pH 7.4, 150 mM NaCl, 5 mM MgCl_2_, 20 mM imidazole, 2% glycerol, 0.2 mg/ml BSA, and 500 μM ATP [(with added [γ-^32^P]-ATP (Hartmann Analytic)]. Prior to addition of phosphatase mix, an aliquot of the reaction was pipetted to SDS sample buffer. Then, a mixture containing Sic1 and phosphatase Cdc14 or PP2A^Cdc55^ was mixed with the phosphorylation reactions to inhibit Clb5-Cdk1 and to measure the dephosphorylation. The final concentration of Sic1 was 1 µM. At 4, 8 and 16 minutes, an aliquot of the reaction was mixed with SDS sample buffer to stop the reaction. The samples were loaded on SDS-PAGE and following electrophoresis, the gels were stained using Coomassie Brilliant Blue R-250 dye and dried.

γ-^32^P phosphorylation signals were detected using an Amersham Typhoon 5 Biomolecular Imager (GE Healthcare Life Sciences). Signals were quantified using ImageQuant TL (Amersham Biosciences). The dephosphorylation assays were performed in three independent replicate experiments.

### Microscope

Images were acquired with Nikon Eclipse Ti2 microscope with 60X Oil CFI Plan APO Lambda lens. Fluorescence illumination is provided by Sola light engine and images were captured by Nikon DS-Qi2. Images were acquired by Nikon NIS element software.

### Time-Lapse movie

Yeast cultures were grown at 30°C in synthetic complete media with 2% glucose (SC) to OD 0.2-0.6, and placed on 2% agarose pad made with SC. Images were acquired at 30°C using a Zeiss Observer Z1 microscope with a 63X/1.4NA oil immersion objective and Orca-r2 C10600-10B camera (Hamatsu Photonics). Images were taken every 3 min for 8 hours. The nuclear fluorescence signals were quantitated using MATLAB (The MathWorks, Inc.). For every strain, data are from at least two repeats with different transformants (Örd *et al*., 2019).

### Protein purification for Mcm2-7 loading assay

Sic1 was purified as described before (Gros *et al*., 2014). ORC and Cdt1·Mcm2-7 were purified as described previously (Remus *et al*., 2009). For Cdc6·Clb2·Cdk1·Cks1, cells (strain YDR12) were grown in 10 L of YPD at 25 °C to a density of 2 x 10^7^ ml^-1^, arrested in G1 phase by addition of 50 μg ml^-1^ α-factor, harvested by centrifugation, resuspended in 10 L YP / 2 % galactose lacking α-factor, and incubated at 25 °C overnight to induce expression of Cdc6-TAP^TCP^. Next day, cells were harvested by centrifugation, washed twice with cold 25 mM Hepes pH 7.6 / 1 M sorbitol, once with buffer H (25 mM Hepes-KOH pH 7.6 / 0.02 % NP-40 / 10 % glycerol) / 300 mM KCl, resuspended in 0.5x volume of buffer H / 300 mM KCl / 2 mM DTT / protease inhibitor cocktail, and frozen as droplets in liquid nitrogen. The resulting ‘popcorn’ was stored at −80 °C until further processing. Cell lysate was prepared by crushing the frozen popcorn in a freezer mill (SPEX CertiPrep 6850 Freezer/Mill) for 6 cycles of 2 minutes at a rate of 15 impacts per second. Crushed cell powder was thawed on ice, resuspended with 1 volume of buffer H / 0.3 M KCl / 1 mM DTT. Insoluble material was pelleted by centrifugation of the lysate in a type 45 Ti rotor (Beckman) for 60 minutes at 40,000 rpm. The clarified extract was supplemented with 2 mM CaCl_2_ and incubated with 1 ml calmodulin affinity beads for 2 hours at 4°C. The calmodulin resin was washed with 10 CV of buffer H / 300 mM KCl / 2 mM CaCl_2_ / 1 mM DTT, and bound protein eluted with 10 CV of buffer H / 300 mM KCl / 1 mM EDTA / 2 mM EGTA / 1 mM DTT. Peak fractions were pooled, concentrated by centrifugation through an Amicon spin concentrator, digested for 4 hours at 4 °C with TEV protease to remove the TAP^TCP^ tag, and subsequently fractionated by gel-filtration using a 24 ml Superdex 200 column equilibrated in buffer H / 300 mM KCl / 1 mM EDTA / 1 mM EGTA / 1 mM DTT. Peak fractions were pooled, diluted with 2x volumes of buffer H / 1 mM EDTA / 1 mM EGTA / 1 mM DTT to a final salt concentration of 100 mM KCl, and fractionated on a 1 ml Mono S ion exchange column using an elution gradient of 0.1 – 1 M KCl over 10 CV. Peak fractions were pooled and stored in aliquots at −80°C after snap freezing in liquid nitrogen. For the original identification of the Cdc6-associated proteins by mass-spectrometry, purified complex was fractionated by SDS-PAGE, individual Coomassie-stained bands excised from the gel, and proteins subjected to mass-spectrometric analysis after in-gel trypsin digestion using standard protocols.

### Gel-filtration analysis of Sic interaction with Cdc6·Clb2·Cdk1·Cks1

6 μg of Sic1, 4 μg Cdc6·Clb2·Cdk1·Cks1, or a mixture of 6 μg Sic1 and 4 μg Cdc6·Clb2·Cdk1·Cks1 were incubated in a 50 μl reaction volume containing buffer H / 300 mM KCl / 1 mM EDTA / 1 mM EGTA / 1 mM DTT for 10 minutes at room temperature, fractionated on a 2.4 ml Superdex 200 PC 3.2/30 column, and fractions analyzed by SDS-PAGE and silver stain.

### Mcm2-7 loading assay

Purified Mcm2-7 loading onto linear 1 kbp ARS-containing DNA immobilized on paramagnetic beads using was performed as described previously (Remus *et al*., 2009), except that 50 nM Cdc6·Clb2·Cdk1·Cks1 was used in place of Cdc6. Where indicated, Sic1 was included at 500 nM.

### Prediction of intrinsically disordered regions in Cdc6 and Clb2

The intrinsically disordered regions and disordered binding regions of Cdc6 were predicted using DISOPRED (Ward *et al*., 2004), IUPRED2A (Mészáros *et al*., 2018), PrDOS, MFDp2 (Mizianty *et al*., 2013), and CSpritz (Walsh *et al*., 2011). A consensus was derived by combining the high confidence predictions from all these programs.

### Structure modeling of Cdc6 and Clb2

The atomic coordinates for Cdc6 isolated from the cryo-EM structure of ORC-Cdc6-Cdt1-Mcm2-7 intermediate (OCCM; PBD ID: 5V8F) (Yuan *et al*., 2017) were extracted and used to build a full-length template of Cdc6. The 140 missing residues from this structure for Cdc6 was modeled separately via an *ab initio* modeling program QUARK (Zhang *et al*., 2016). The full-length Cdc6 model was constructed by combining above mentioned fragment and extracted coordinates of Cdc6 using MODELLER (Eswar *et al*., 2006) and further refined using 3DRefine (Bhattacharya *et al*., 2016). Clb2 was modeled using a hybrid threading approach, *ab initio*, and template-based approach based on the algorithm, I-TASSER (Iterative Threading ASSEmbly Refinement) (Roy *et al*., 2010). I-TASSER identifies structural templates from the PDB by a multi-threading server, LOMETS (Wu & Zhang, 2007), and then constructs full-length atomic models by iterative template-based fragment assembly simulations. The IDR in Clb2 was modeled independently using QUARK (Zhang *et al*., 2016) and incorporated to create a full-length model in similar manas described above for Cdc6. Both the full-length modeled structures were meticulously evaluated by various structure verification programs, including VERIFY3D (Eisenberg *et al*., 1997), Voromqa (Olechnovič & Venclovas, 2017), Proq3 (Uziela *et al*., 2016), and ProsaWeb (Wiederstein & Sippl, 2007). These programs employ a variety of methods inclusive of deep learning approaches to evaluate the quality of the models. The final models were chosen by selecting the best evaluation profiles and their correlation with known functional features. The models were visualized and analyzed for their biophysical properties using the visualization programs Pymol (Schrodinger, 2010), and ChimeraX (Pettersen *et al*., 2021).

## Acknowledgement

We would like to thank Lea Schroeder and Yooko Caroll for technical assistance, and Dr. Frederick Cross for anti-Clb2 antibody. This work was supported by PSC-CUNY enhanced award to A.I., NIGMS grants R01-GM127428 and R01-GM107239 and NIH/NCI Cancer Center Support Grant P30 CA008748 to D.R, ERC Consolidator Grant 649124, Centre of Excellence for “Molecular Cell Technologies” TK143, and Estonian Science Agency grant PRG550 to M. L, Francis Crick Institute, which receives its core funding from Cancer Research UK, the UK Medical Research Council, and the Wellcome Trust (FC001066) to J.D.

---

**Table S1.** Strain List

| Strain# | Genotypes | Origin |
| --- | --- | --- |
| JPY003 | *MATa bar1 CDC6-9MYC::TRP1 ADE2 leu2-3 trp1-1 ura3-1 his3-11* | This study |
| JPY009 | *MATa bar1 Δclb2::ADE2 CDC6-9MYC::TRP1 ade2-1 leu2-3 trp1-1 ura3-1 his3-11* | This study |
| JPY011 | *MATa bar1 Δcdc55::KanMX CDC6-9MYC::TRP1 ADE2 leu2-3 trp1-1 ura3-1 his3-11* | This study |
| JPY005 | *MATalpha CDC6-9MYC::TRP1 Δclb5::URA3 ade2-1 leu2-3 trp1-1 ura3-1 his3-11* | This study |
| JPY002 | *MATa bar1 CDC6-9MYC::TRP1 Δrts1::KanMX ADE2 leu2-3 trp1-1 ura3-1 his3-11* | This study |
| JPY029 | *MATa bar 1 CDC6-prA::HIS3 ade2-1 leu2-3 trp1-1 ura3-1 his3-11* | This study |
| JPY030 | *MATa bar 1 CDC6-prA::HIS3 Δclb2::LEU2 ade2-1 leu2-3 trp1-1 ura3-1 his3-11* | This study |
| JPY032 | *MATa bar 1 CDC6-PRA::HIS CDC55-MYC::KanMX ade2-1 leu2-3 trp1-1 ura3-1 his3-11* | This study |
| JPY033 | *MATa bar1 CDC6-PRA::HIS cdc55-101-MYC::KanMX ade2-1 leu2-3 trp1-1 ura3-1 his3-11* | This study |
| JPY183 | *MATalpha whi5::Whi5-mCherry::SpHIS5 ura3::PCDC6-CDC6-yeCitrine::URA3 cdc6::natNT2::GALS-3HA-CDC6 ADE2 leu2-3 trp1-1 ura3-1 his3-11* | from Mart Loog |
| JPY175 | *MATalpha cdc55-101-MYC::KanMX whi5::Whi5-mCherry::SpHIS5 ura3::PCDC6-CDC6-yeCitrine::URA3 cdc6::natNT2::GALS-3HA-CDC6 ADE2 leu2-3 trp1-1 ura3-1 his3-11* | This study |
| JPY174 | *MATalpha Δcdc55::KanMX whi5::Whi5-mCherry::SpHIS5 ura3::PCDC6-CDC6-yeCitrine::URA3 cdc6::natNT2::GALS-3HA-CDC6 ADE2 leu2-3 trp1-1 ura3-1 his3-11* | This study |
| JPY193 | *MATalpha CDC6-GFP::HIS3 ADE2 leu2-3 trp1-1 ura3-1 his3-11* | This study |
| JPY192 | *MATalpha cdc55-101-MYC::KanMX CDC6-GFP::HIS3 ADE2 leu2-3 trp1-1 ura3-1 his3-11* | This study |
| JPY056 | *MATa bar 1 CDC6-T368A-PRA::HIS3 ade2-1 leu2-3 trp1-1 ura3-1 his3-11* | This study |
| JPY057 | *MATa bar 1 CDC6-T368A-PRA::HIS3 Δclb2::LEU2 ade2-1 leu2-3 trp1-1 ura3-1 his3-11* | This study |
| JPY048 | *MATalpha Δclb2::LEU2 ade2-1 leu2-3 trp1-1 ura3-1 his3-11* | This study |
| BCY581 | *MATa bar1 CDC6-T39A-T368A-prA::HIS3 ade2-1 leu2-3 trp1-1 ura3-1 his3-11* | This study |
| JPY123 | *MATa bar 1 URA3::CDC6-T7A-T23A-T39A-S43A-T368A-S372A-13MYC ADE2 leu2-3 trp1-1 his3-11* | This study |
| JPY126 | *MATalpha Δcdc55::KanMX URA3::CDC6-T7A-T23A-T39A-S43A-T368A-S372A-13MYC ade2-1 leu2-3 trp1-1 his3-11* | This study |
| JPY124 | *MATa bar 1 URA3::CDC6-T23A-T39A-S43A-T368A-S372A-13MYC ADE2 leu2-3 trp1-1 his3-11* | This study |
| JPY127 | *MATa bar 1 Δcdc55::KanMX URA3::CDC6-T23A-T39A-S43A-T368A-S372A-13MYC ade2-1 leu2-3 trp1-1 his3-11* | This study |
| JPY087 | *MATalpha URA3::GAL-SIC1-HA CDC6-T39A-T368A-prA::HIS3*<br>*Δcdh1::LEU2 ade2-1 leu2-3 trp1-1 his3-11* | This study |
| JPY069 | *MATalpha URA3::GAL-CDC6-HA ADE2 leu2-3 trp1-1 his3-11* | This study |
| JPY066 | *MATalpha Δclb2::LEU2 URA3::GAL-CDC6-HA ADE2 leu2-3 trp1-1*<br>*his3-11* | This study |
| JPY068 | *MATa URA3::GAL-CDC6-T368A-HA ADE2 leu2-3 trp1-1 his3-11* | This study |
| JPY065 | *MATalpha Δclb2::LEU2 URA3::GAL-CDC6-T368A-HA ADE2 leu2-3*<br>*trp1-1 his3-11* | This study |
| JPY067 | *MATalpha URA3::GAL-CDC6-T39A-T368A-HA leu2-3 trp1-1 his3-11* | This study |
| JPY064 | *MATalpha Δclb2::LEU2 URA3::GAL-CDC6-T39A-T368A-HA leu2-3*<br>*trp1-1 his3-11* | This study |
| JPY117 | *MATa bar 1 CDC6-9MYC::TRP1 cdc14-3 ade2-1 leu2-3 trp1-1 ura3-1*<br>*his3-11* | This study |
| JPY129 | *MATa bar 1 CDC6-9MYC::TRP1 cdc15-1 ade2-1 leu2-3 trp1-1 ura3-1*<br>*his3-11* | This study |

