## Supplemental Figures for "Cdc6 is sequentially regulated by PP2A-Cdc55, Cdc14 and Sic1 for origin licensing in *S. cerevisiae*"

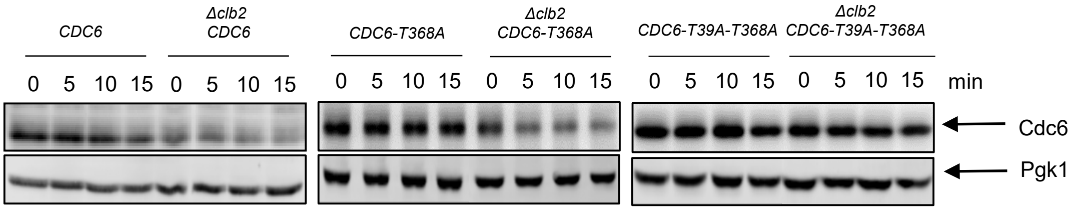


Figure S1. Cdc6 suppression in *Δclb2* is through Cdc6 N-terminal phospho-degron. A) *GAL-CDC6-PrA (CDC6), GAL-CDC6-T368A-PrA (CDC6-T368A), GAL-CDC6-T39A-T368A-PrA (CDC6-T39A-T368A), Δclb2 GAL-CDC6-PrA (Δclb2 GAL-CDC6), Δclb2 GAL-CDC6-T368A-PrA (Δclb2 GAL-CDC6-T368A),* and *Δclb2 GAL-CDC6-T39A-T368A-PrA (Δclb2 GAL-CDC6-T39A-T368A)* cells were in blocked in mitosis by nocodazole. Galactose was added to induce Cdc6 overexpression. Glucose was added at time 0 to repress transcription, and Cdc6 degradation was measured for 15 minutes. Protein was extracted and subjected to western blot analysis to visualize Cdc6-PrA. Pgk1 was used as a loading control.

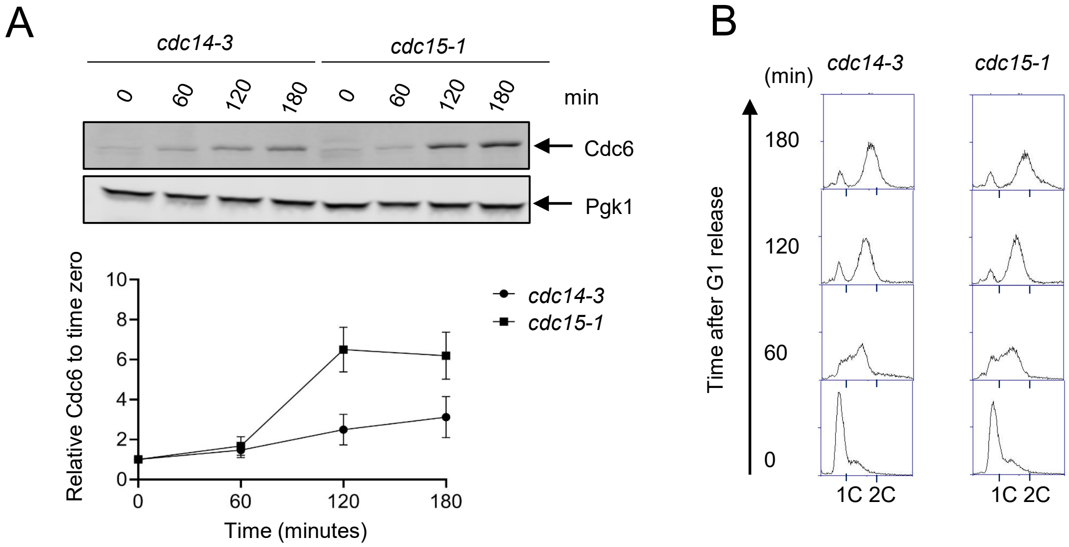


Figure S2. Cdc6 is unstable in *cdc14-3* mutant. A) *CDC6-9MYC cdc14-3* (*cdc14-3*) or *CDC6-9MYC cdc15-1* (*cdc15-1*) cells were synchronized in G1 phase by alpha-factor first. The cell cycle was released and samples were collected every 60 minutes up to 180 minutes at 37 degrees. Western blot analysis was performed to visualize Cdc6-MYC. Pgk1 was used as a loading control. The relative Cdc6 band intensity to time zero from is shown. Error bar is SEM (n=3). B) Samples from A were fixed and stained with PI to show cell cycle profiles by flow cytometry.

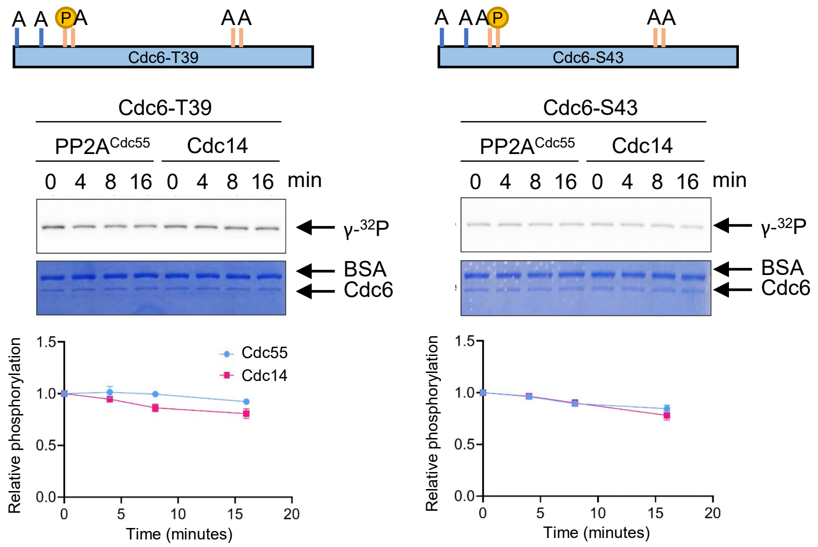


Figure S3. PP2A and Cdc14 do not target Cdc6-T39 and S43. Cdc6-T7A-T23A-S43A-T368A-S372A (Cdc6-T39) or Cdc6-T7A-T23A-T39A-T368A-S372A (Cdc6-S43) was phosphorylated by purified Clb5-Cdk1 in the presence of [γ-$^{32}$P]-ATP. Phosphatase activity of Cdc14 or PP2A$^{Cdc55}$ towards these proteins was measured as described in Figure 4.

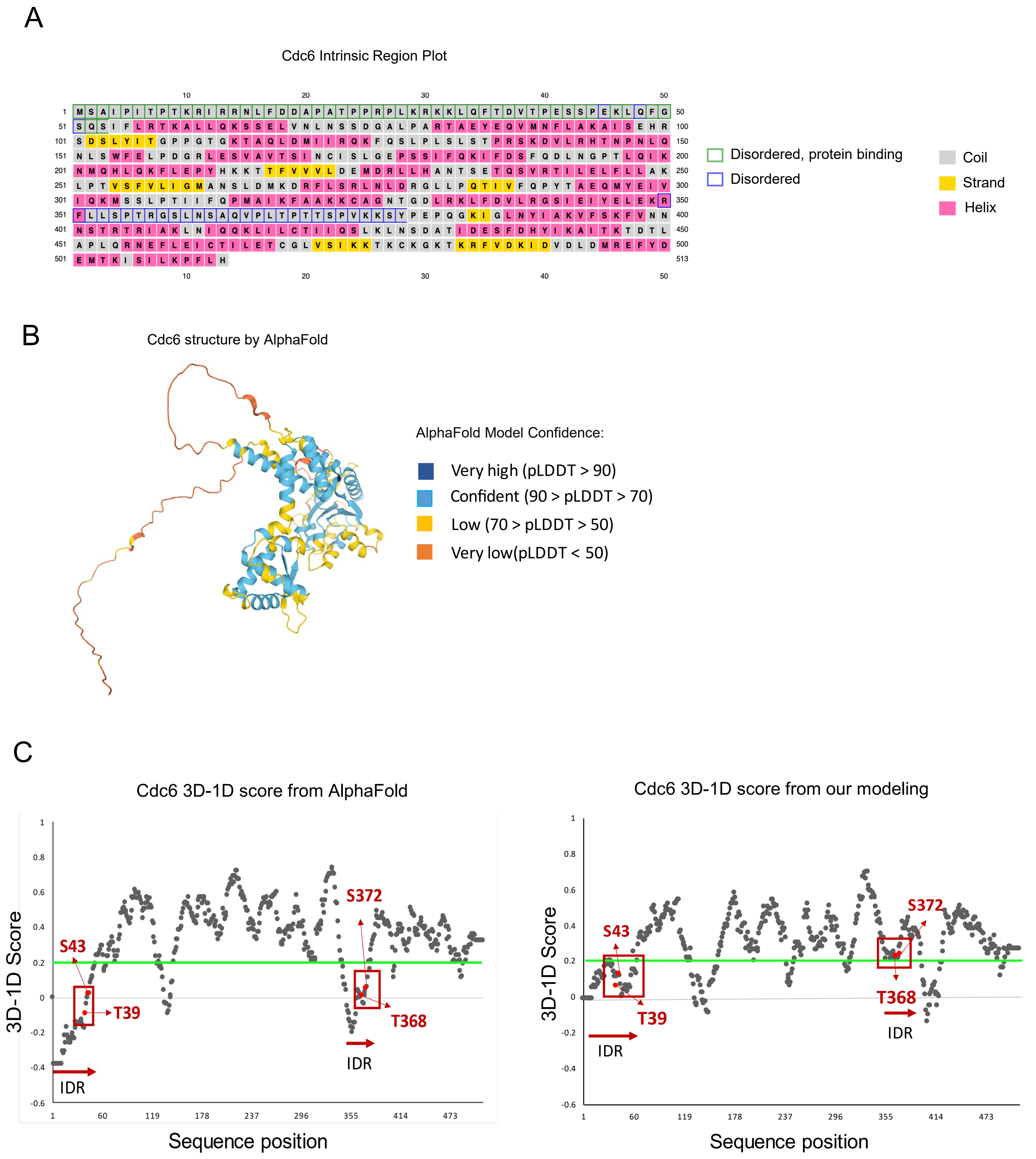


Figure S4. A prediction of Intrinsically Disordered Region (IDR) in Cdc6. A) Cdc6 IDR predictions by DISOPRED and secondary structure prediction by IUpred2. B) Cdc6 structure from AlphaFold with internally generated confidence scores. C) 3D-1D Score by Verify3D program are shown for Cdc6 structures by AlphaFold and our modeling program. The green line represents the 3D-1D score = 0.2 which indicates threshold for model quality. The red boxes display the location of phospho-degrons (T39-S43;T368-S372). Each amino acid residue is distinguished by arrows and the red horizontal arrows indicate the position of IDRs.
